# Olfactory Loss Enhances Visual Learning in *Drosophila* through Structural and Functional Reorganisation

**DOI:** 10.64898/2026.06.16.732444

**Authors:** Büşra Çoban, Samuel N. Harris, Burak Gür, Marc Corrales, Ana Jesus Correia Da Silva, Diana Shevchuk, Kerstin Leptien, Dennis Goldschmidt, Christopher Schnaitmann, Anissa Kempf, Michał Januszewski, Nicolo Ceffa, Michael Clayton, Ivana Henry, Shi Yan Lee, Amina Dulac, Albert Cardona, Marta Zlatic, Johannes Felsenberg

## Abstract

Loss of a sensory modality can enhance performance in the remaining senses. However, the circuit mechanisms by which such cross-modal compensation can improve cognitive functions, including learning, are unknown. Here, we show that compromising olfaction in both larval and adult *Drosophila* enhances visual associative learning. Using behavioural analysis, functional imaging, and comparative connectomics, we reveal the circuit mechanisms that underlie this improvement. Animals with improved learning ability have enhanced responses to visual stimuli in the higher-order learning circuit. The complementary circuit mechanisms that can enhance these responses are structural reweighting of inputs in the larva, resulting in an increased fraction of synaptic inputs from visual pathways onto neurons in the learning circuit, and a reduction in cross-modal inhibition in the adult. Together, these findings reveal synaptic and disinhibitory circuit mechanisms that enhance learning in higher-order associative networks following sensory loss.

## Introduction

To survive, animals rely on extracting and integrating salient sensory information from their surroundings. Loss of a sensory modality can trigger a reorganisation of neural circuits in the brain, often accompanied by enhanced behavioural responses to the remaining sensory systems (Bavelier & Neville, 2002; Lee & Whitt, 2015; López-Bendito et al., 2022; Merabet & Pascual-Leone, 2010). For example, blindness in humans can lead to superior auditory localisation (Alary et al., 2008; Cuevas et al., 2009; Lessard et al., 1998; Röder et al., 1999), mosquitoes (*Aedes aegypti*) with impaired olfaction rely more strongly on thermosensation to locate hosts (Morita et al., 2025), and nematode worms (*Caenorhabditis elegans*) lacking mechanosensation exhibit enhanced chemotaxis (Rabinowitch et al., 2016). These adaptive neuronal processes are collectively referred to as cross-modal plasticity (Bavelier & Neville, 2002; Lee & Whitt, 2015; Merabet & Pascual-Leone, 2010). In general, two broad changes can occur following sensory loss: the deprived primary sensory pathway can become responsive to input from other modalities (cross-modal recruitment), while brain areas processing the remaining sensory modalities show refined activity (cross-modal compensation) (Ewall et al., 2021; Goel et al., 2006; Jitsuki et al., 2011; Merabet & Pascual-Leone, 2010). However, while cross-modal compensation can influence behaviour and higher-order processing (Amedi et al., 2003; Raz et al., 2007), the circuit mechanisms underlying its effects on learning and memory remain poorly understood.

Mechanistic insight into cross-modal plasticity in associative centres requires access to identified circuit components that mediate memory-related functions, enabling analysis of synaptic activity and connectivity at cellular resolution. Decades of research in the fruit fly, *Drosophila melanogaster*, have established a detailed understanding of the neuronal circuitry underlying associative learning in both larval and adult stages (Cognigni et al., 2018; Eschbach & Zlatic, 2020; Thum & Gerber, 2019). Adult and larval flies can learn to associate olfactory or visual cues with reward or punishment (Aceves-Piña & Quinn, 1979; Gerber et al., 2004; Quinn et al., 1974; Quinn & Dudai, 1976; Scherer et al., 2003; Tempel et al., 1983). These memories are stored in the mushroom body (MB) as dopamine-dependent synaptic modifications between stimulus-encoding Kenyon cells (KCs) and specific mushroom body output neurons (MBONs), which bias behaviour toward approach or avoidance (Aso et al., 2010; Aso, Sitaraman, et al., 2014; Burke et al., 2012; Eichler et al., 2017; Eschbach et al., 2020; Gerber et al., 2004; Liu et al., 2012; Owald et al., 2015; Saumweber et al., 2018; Séjourné et al., 2011; Thum & Gerber, 2019; Vogt et al., 2014). Distinct KC subtypes receive modality-biased input: most KCs receive input from olfactory projection neurons, whereas smaller subsets receive visual input directly or indirectly from the optic lobes (Aso, Hattori, et al., 2014; Crittenden et al., 1998; Eichler et al., 2017; Ganguly et al., 2024; Li et al., 2020; Strausfeld et al., 2003; Vogt et al., 2016; Yagi et al., 2016; Zheng et al., 2018). These input pathways have been mapped at synaptic resolution in whole-brain connectomes for both larvae and adults (Eichler et al., 2017; Li et al., 2020; Takemura et al., 2017; Winding et al., 2023; Zheng et al., 2018). The multimodal architecture of the MB makes it a compelling candidate substrate for cross-modal compensation following sensory loss.

Cross-modal plasticity depends on interactions between the deprived and the intact sensory pathways. Theories describing this interaction broadly fall into three categories: the formation of new ectopic connections, synaptic reweighting through intrinsic cellular or homeostatic mechanisms, and a flexible rebalancing of activity through existing circuit architecture including disinhibitory motifs (Barnes et al., 2017; Bridi et al., 2018; Ewall et al., 2021; Keck et al., 2017; Kral & Sharma, 2023; Lee, 2023; Tripodi et al., 2008). Early studies in mammals suggested that perturbations to sensory pathways precipitate large-scale rewiring of sensory brain areas (Sur et al., 1990; von Melchner et al., 2000). However, more recent evidence indicates that large-scale reorganisation is not required for sensory enhancement; instead, compensation often arises from modulation of pre-existing multisensory connections and inhibitory circuits (Iurilli et al., 2012; Kral et al., 2005; Kral & Sharma, 2023; Lee, 2023; Makin & Krakauer, 2023; Petrus et al., 2014, 2015; Rabinowitch et al., 2016; Whitt et al., 2022). For example, disinhibitory circuit motifs have been proposed to mediate activity upregulation following sensory deprivation (Whitt et al., 2022), and homeostatic adjustments can strengthen previously subthreshold intermodal inputs (Ewall et al., 2021; Kral & Sharma, 2023; Lee & Whitt, 2015; Valdes-Aleman et al., 2021). However, it is unknown whether similar principles could operate within higher-order learning circuits (such as the MB) to improve learning with the remaining sensory modalities following loss of one modality. Light-level analyses suggest that olfactory loss does not induce gross rewiring of MB inputs (Hayashi et al., 2022), but whether it improves visual learning and triggers synaptic-level reorganisation, altered inhibitory balance, or recruitment of latent pathways has not been tested.

Here, we leverage the experimental tractability of *Drosophila* to determine whether and how loss of olfactory input enhances visual memory. By genetically and surgically perturbing olfaction, we show that this deprivation enhances visual learning in both larvae and adults. Using comparative connectomics and functional analyses, we uncover complementary circuit adaptations that converge on elevated visually-driven activity in the MB. These findings provide mechanistic insight into how higher-order associative circuits reorganise to promote cognitive flexibility and improve learning.

## Results

### Olfactory deprivation enhances visual learning in flies

Loss of one sensory modality can improve performance in others, but whether such compensation extends to associative memory is less clear. We therefore tested whether compromising olfactory function alters visual associative memory in both larval and adult *Drosophila*. Because odour detection requires the olfactory co-receptor Orco, we first asked whether visual learning is altered in animals with the *orco* loss-of-function allele o*rco¹ (Fishilevich et al., 2005; Larsson et al., 2004)*.

To examine visual associative learning in larvae, we adapted a previously reported quantitative, high-throughput learning assay for olfactory conditioning (Croteau-Chonka et al., 2022). During training, the conditioned stimulus (CS; green light) was paired with an unconditioned stimulus (US; intense heat) in either forward or backward temporal order (Fig. 1A). Stimulus valence was estimated using a ratio of turning responses at the onset and offset of stimulation in line with previous studies (Croteau-Chonka et al., 2022; Gepner et al., 2015; Gomez-Marin et al., 2011; Klein et al., 2017), which were quantified as a behavioural preference index (BPI, see Methods). To account for non-associative effects, memory acquisition was quantified as the difference in BPI in the testing trials between forward-paired and backward-paired groups of larvae, which we refer to as the behavioural learning index (BLI). We trained groups of Canton-S (wild-type) and o*rco¹* larvae (*orco* mutant), raised to the L3 feeding stage, with either the forward-paired or backward-paired protocol and then tested their turning responses to visual stimulation. Tests of olfactory preference confirmed that *orco* mutant larvae were olfactorily deprived (Fig. S1A). For both *orco* mutant and wild-type larvae, forward-paired training produced significantly more aversive turning responses than backward-paired training, demonstrating that both genotypes can form visual memory (Fig. 1B, Fig. S1B). Furthermore, we found that *orco* mutant larvae exhibit significantly stronger aversive visual memory than wild-type controls (Fig. 1C). Similar effects were observed at the L2 stage (Fig. S1C-D), indicating that enhanced visual learning in olfactory-compromised animals begins early in larval development.

**Figure 1:**
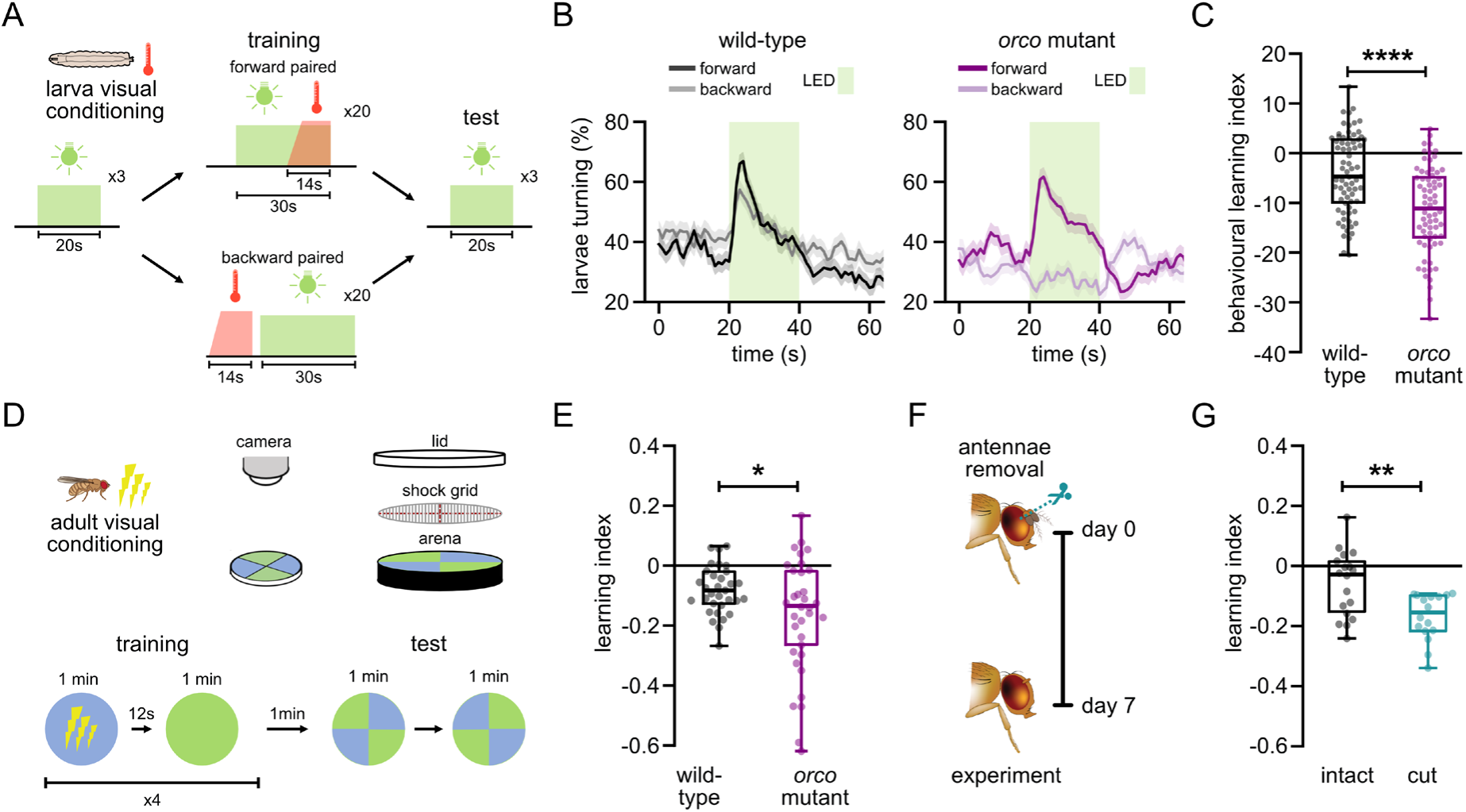
Sensory deprivation enhances visual learning. A) Visual conditioning protocol for larval flies. Turning behaviour was recorded for larvae that underwent either forward-paired (CS followed by US) or backward-paired (US followed by CS) training protocols. Afterwards, the learned behaviour was tested. B) Comparison of larval turning during the test quantified as a behavioural preference index (see Methods) between wild-type (Canton-S, grey) and orco mutants (orco^1^, magenta). Traces show mean ± SEM. C) Behavioural learning index quantified from the turning differences between forward- and backward-trained wild-type (grey) and orco mutant (magenta) larvae. ****p<0.0001, Mann-Whitney U test. D) Visual conditioning paradigm in adult flies. Groups of flies undergo a training period with a specific colour (CS+, either blue or green) paired with shock (US), followed by a test period where the learned behaviour is assessed. E) Quantification of learning index based on the change in colour preferences of the flies between training and testing periods. Comparison of wild-type (grey) and orcomutants (magenta). *p<0.05, two-tailed Student’s t-test. F) Antennae of adult flies were removed after eclosion (day 0) and experiments were performed on day 7. G) Comparison of the learning index between flies with intact antennae (grey) and flies that had their antennae cut (teal). **p<0.05, two-tailed Student’s t-test. Boxplots in (C, E, G) show median, interquartile ranges and whiskers extend to the most extreme data points. Sample sizes for B-C) wild-type: forward n=71, backward n=72, orco mutant: forward n=68, backward n=63 (larvae); (E) wild-type: n= 32, orco mutant: n= 34 (groups of flies); (G) intact: n=18, cut: n=16 (groups of flies).

To test whether the enhancement observed in larvae also occurs in adults, we examined visual learning in seven-day-old adult wild-type and age-matched *orco* mutants (*orco^1^*). Following previously established protocols in adults (Schnaitmann et al., 2013; Vogt et al., 2014), we assayed visual learning in custom-built circular training arenas (Fig. 1D). During training, groups of flies were exposed to 1-minute illumination with either blue or green light (paired CS, CS+) paired with 120 V electric shocks (US), followed by a 12-second interval before presenting the other colour without reinforcement (unpaired CS, CS-). This training cycle was repeated four times, and memory was tested by allowing flies to choose between the previously shock-paired CS+ and the unpaired CS-displayed in opposing quadrants of the arenas (Fig. 1D). Learning performance was averaged across two reciprocal trainings in which each colour served as the CS+, controlling for training-unrelated biases. As in larvae, adult *orco* mutants displayed enhanced visual learning compared to wild-type controls (Fig. 1E). Together, these results indicate that compromise of olfactory function can increase visual memory capacity across paradigms and developmental stages.

We next asked whether enhanced visual learning is also evident following more acute sensory loss. To this end, we surgically removed the primary olfactory organs (antennae) in adult flies within 24 h of eclosion. Flies with cut antennae initially showed an increase in daytime sleep, indicating ongoing recovery from injury (Fig. S1E) (Singh & Donlea, 2020), yet sleep levels returned to baseline at day 7, suggesting the completion of recovery (Fig. S1F). Thus, we performed experiments 7 days after antennae removal (Fig. 1F). Cutting the antennae abolished flies’ innate avoidance of aversive odours, indicating perturbation of olfaction (Fig. S1G). In visual conditioning, intact controls showed moderate learned avoidance of the shock-paired colour. Strikingly, relative to intact controls, antennal removal enhanced learned avoidance (Fig. 1G). Importantly, when repeating the experiments under a mock condition (no electric shock during training) learning was absent in both groups (Fig. S1H). Together, these data indicate that sensory deprivation by acute loss of the primary olfactory organs enhances adult visual memory performance.

Collectively, using different visual associative memory paradigms in larval and adult flies revealed a consistent cross-modal memory enhancement: perturbing olfactory input improves visual associative learning. Because neural substrates of associative memory are well mapped in the fly brain (Cognigni et al., 2018; Eschbach & Zlatic, 2020; Thum & Gerber, 2019), these results provide a framework to leverage the strengths of the larval and adult systems to address where and how this enhancement is implemented.

### Olfactory deprivation increases visual synaptic input to neurons in the larval learning circuit and enhances their responses to visual stimuli

To uncover the circuit mechanisms that could underlie improved visual learning, we first asked whether synaptic input from visual pathways onto MB neurons is increased in the absence of olfaction. While the greatest number of sensory inputs to the MB in *Drosophila* are olfactory, anatomical and functional studies in larvae and adults show that visual signals also reach KCs (Eichler et al., 2017; Larderet et al., 2017; Li et al., 2020; Yagi et al., 2016).

In adults, visual input to the MB is spatially segregated, with approximately 8% of all KCs receiving predominantly visual input at specialized accessory calyces (Ganguly et al., 2024; Li et al., 2020). This visual processing is mediated by two distinct KC populations: γd-KC cells that receive segregated input at the ventral accessory calyx (vACA) and KCαβp cells that receive input at the dorsal accessory calyx (dACA).

In the larval system, photoreceptor neurons relay visual information from the Bolwig organ to nine visual projection neurons (vPNs) that form distinct pathways to central targets (Humberg & Sprecher, 2017; Keene et al., 2011; Larderet et al., 2017; Malpel et al., 2002). While four projection neurons contribute to circadian entrainment (Kaneko et al., 1997; Malpel et al., 2002), four vPNs (vPN1-4) receive specialized input and project toward the lateral horn and regions adjacent to the MB calyx (Larderet et al., 2017).

We analysed the larval whole-brain connectome and confirmed previous findings that only a few KCs receive direct visual input, highlighting the predominance of olfactory pathways (Fig. S2A) (Eichler et al., 2017; Larderet et al., 2017; Winding et al., 2023). Two KCs per brain hemisphere receive significant (>3% axodendritic) inputs from vPNs, which we term the Larval Optic Neuropil KC (*lon*KC) and the olfactory-visual KC (*ov*KC). These KCs integrate multimodal information, with both receiving inputs from vPNs and multiglomerular olfactory PNs (oPNs, Fig. S2B). These cells differ from classical olfactory KCs by lacking defined dendritic claws, a characteristic also observed in visual adult Kenyon cell types (Li et al., 2020), and projecting dendrites to regions posterior and ventral to the MB calyx (Fig. S2C). These multisensory KCs may serve as a hub for cross-modal integration, potentially facilitating cross-modal compensation that enhances visual learning upon the loss of olfaction.

To investigate whether a structural rebalancing of these key sites of multisensory integration occurs following the loss of olfactory signalling, we imaged the entire brains of an L2-stage anosmic mutant (*orco^1^)* and a wild-type control larva using enhanced focused ion-beam-scanning electron microscopy (eFIB-SEM) with 8 x 8 x 8 nm resolution (Fig. 2A-B). We chose to do this analysis in the L2 stage, as the L3 stage and the adult brain are too big to be imaged in a reasonable amount of time. After imaging the larval brains, we automatically segmented neuronal projections, as previously described (Januszewski et al., 2018; Scheffer et al., 2020; Xu et al., 2017). We identified *ov*KC and *lon*KC based on their characteristic dendritic projections, proof-read them, annotated their synaptic connections and then proof-read their strongest (>3% axodendritic input in both hemispheres) upstream partners. These strongest partners accounted for over 70% of the total axodendritic input onto *ov*KC and *lon*KC in both volumes (wild-type: *ov*KC - 72.8%, *lon*KC - 74.7%; *orco^1^*: *ov*KC - 73.2%, *lon*KC - 71.7%). Based on comparisons to the L1-stage connectome, we confidently and uniquely identified all of these strong partners with two exceptions: we could not distinguish between the two local neurons of the BVLA12 lineage (referred to as bvla12L), nor could we distinguish the ipsilateral and contralateral projections of oPN2 from one another. In both of these cases their outputs onto KCs were summed to allow for comparison between the brains. In both the anosmic (*orco* mutant) and the wild-type brain, vPN1 synapses onto *ov*KC, while vPN2 and vPN3 connected to *lon*KC, recapitulating the organisation observed in the L1-stage connectome (Fig. 2C, 2D) (Winding et al., 2023). Furthermore, the connections from oPNs, interneurons and local neurons were highly conserved between both brains, suggesting that these multisensory KCs receive a stereotyped set of inputs. We then compared the percentage input of homologous KC input neurons between the two volumes. The average fraction of input varied by 1.12 ± 0.18 (fold-change, mean ± SEM) for the 18 homologous neurons we identified.

**Figure 2:**
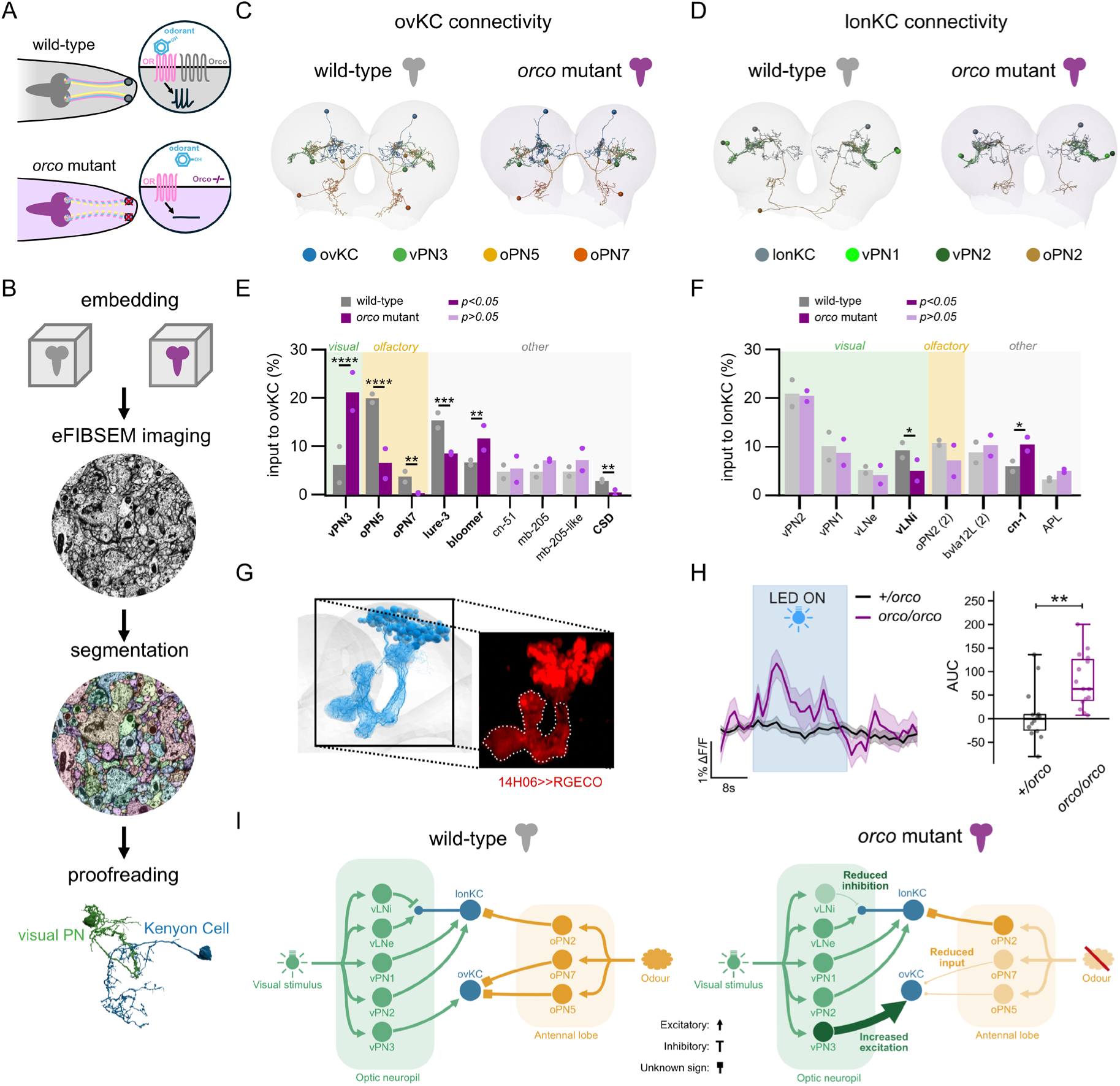
Loss of olfactory signalling is associated with strengthening of visual projection neurons to Kenyon cell connections. A) Schematic showing wild-type larva (grey), where Orco association with the olfactory receptors (OR) leads to increase in downstream neural activity upon odorant binding. In orco mutants (magenta), the lack of Orco prevents odorant-based neural activation. B) Schematic showing the steps used for imaging the brains of an L2 orco mutant (orco^1^, magenta) and a wild-type (Canton-S, grey) larva. C, D) The skeletons of multisensory Kenyon cells, ovKC (blue, C) and lonKC (slate, D) and their major inputs: olfactory PNs oPN5 (yellow), oPN7 (orange), and oPN2 (gold) and visual PNs vPN3 (green), vPN1 (light green), and vPN2 (dark green) in the wild-type (grey) and orco mutant (magenta) volumes. E, F) The fraction of synaptic input from all KC upstream partners with greater than 3% bilateral input in either the orco mutant or wild-type volume onto ovKC (E) and lonKC (F). Inputs from the ipsilateral and contralateral projections of oPN2 were summed, as well as the inputs from two local neurons of the BVLA12 lineage (denoted by ‘(2)’). *p<0.05, **p<0.01, ***p<0.001, ****p<0.0001, chi-square test followed by Benjamini-Hochberg correction. Sample sizes: one individual larva for L2 wild-type (Canton S) and one individual larva for L2 orco mutant (orco^1^). G) Electron microscopy reconstruction of KCs from the L1-connectome (left) and a projection of the 3D light microscopy volume showing the MB lobes in outline (right). H) Calcium responses of KCs in the MB lobes upon visual stimulation for homozygous orco mutant larvae (purple) and heterozygous control larvae (black). Comparison of the response area under the curve (AUC) between genotypes. **p<0.01 Mann-Whitney U Test. Sample sizes: orco/orco = 12, +/orco = 11. I) Schematic connectivity diagram highlighting the structural changes observed in visual and olfactory neurons between the wild-type (grey) and orco mutant (magenta) volumes. Connections previously shown to be from cholinergic neurons were assumed to be excitatory whilst glutamatergic connections were assumed to be inhibitory.

Quantitative analysis of the percentage of synaptic input from projection neurons onto *ov*KC revealed three significant changes in structural connectivity: the connection from the vPN3 was greatly strengthened (3.3-fold increase) in the anosmic (*orco* mutant) animal relative to the wild-type, while inputs from two distinct multiglomerular olfactory PNs, oPN2 and oPN7, were strongly weakened, as well as from the modulatory CSD neurons that also received input from olfactory sensory neurons (Fig. 2E, Fig. S2D). We also saw significant changes in two other neurons that did not receive direct sensory neuron input and whose function is unknown, making it harder to interpret the consequences of these changes (lure-3, bloomer, Fig. 2E). For all other strong upstream partners of *ov*KC no significant variation in input fraction was identified (Fig. 2E). Therefore, our findings suggest a broad reweighting of connections onto *ov*KC with visual inputs strengthening at the expense of olfactory input within *orco* mutants.

In contrast, no significant changes in the fractions of synaptic input from visual (vPN1 and vPN2) or olfactory (oPN2) projection neurons onto *lon*KC were observed between the anosmic (*orco* mutant) and the wild-type brain (Fig. 2G and S2E). *lon*KC is unique in that it projects a long dendrite directly into the primary visual sensory region, the larval optic neuropil (LON), and forms synapses with a range of visually sensitive neurons therein, including two local neurons: the excitatory vLNe and the inhibitory vLNi (Daniels et al., 2008; Larderet et al., 2017). Interestingly, we observed a reduction in the fraction of synaptic input from the vLNi in the anosmic larva, compared to wild-type (Fig. 2F). Therefore, *lon*KC may undergo a different structural compensatory change: namely a reduction in inhibitory synaptic input from visual pathways (rather than an increase in excitatory input). In addition, we observed a significant increase in connectivity from the interneuron, cn-1, that does not receive sensory neuron input and whose function is unknown, making it difficult to interpret the consequence of the change. Overall, the larval connectomic analysis suggests that loss of olfactory input increases the fraction of visual excitatory synaptic input to the MB through vPN3 onto the multisensory *ov*KC and decreases the fraction of inhibitory visual synaptic input onto the *lon*KC (Fig. 2I). Previously, it has been shown that an increase in the fraction of excitatory synaptic input from a presynaptic partner leads to enhanced responses to that partner (Valdes-Aleman et al., 2021). We, therefore, wanted to confirm whether KCs in anosmic (*orco* mutant) larvae have enhanced responses to visual stimuli by imaging their activity. Selective GAL4 lines for targeting individual KC subtypes are not available in the larva, so we used a GAL4-line (14H06-Gal4, Jenett et al., 2012) to target a calcium indicator of neural activity (RGECO, Dana et al., 2016) to all KCs and imaged average calcium signals of all KC axons in the MB lobes of L2 larvae (Fig. 2G). We found that the mean response of KCs to visual stimuli was significantly increased in anosmic (*orco* mutant) compared to heterozygous control larvae (Fig. 2H). Taken together, our findings are consistent with the idea that increased excitatory synaptic input from visual pathways onto the multisensory *ov*KC and decreased inhibitory visual synaptic input onto the *lon*KC lead to enhanced responses of these KCs to visual stimuli which could, in turn, contribute to enhancing visual learning (Fig. 2I).

### Olfactory deprivation enhances activity of neurons that receive visual input in the adult learning circuit and recruits an additional subtype of visual KCs to the memory circuit

Next, we wanted to examine whether olfactory deprivation also enhances activity of neurons that receive visual input in the adult learning circuit. In the adult fly, KCs can be divided into seven subtypes based on anatomy and connectivity (Aso, Hattori, et al., 2014). While most subtypes receive predominantly olfactory input, two populations, the γd-KCs and the αβp-KCs, receive direct visual input (Fig. 3A) (Li et al., 2020; Yagi et al., 2016).

**Figure 3:**
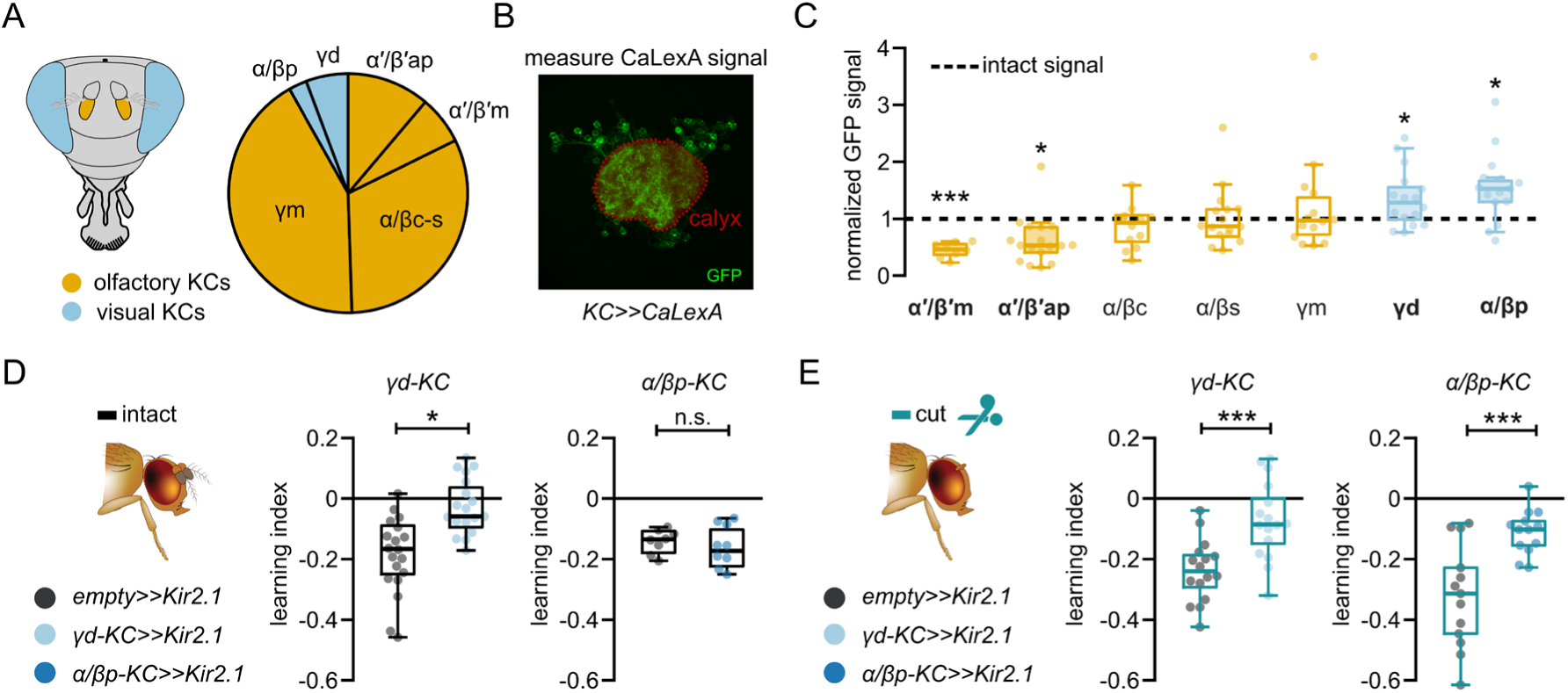
Olfactory loss shifts activity in the MB and recruits a specific KC subtype into memory circuitry. A) Overview of the proportion of olfactory (dark yellow) and visual KCs (blue) based on the FAFB/FlyWire connectome. B) Assessment of KC activity based on CaLexA signals in the calyx region. C) Comparison of GFP signals of different KC subtypes in cut adult flies, each normalized to the GFP signals in respective controls of intact adult flies. Sensory deprivation increased activity in visual KCs (blue) and some olfactory KCs (dark yellow) had decreased activity. *p<0.05, ***p<0.001, two-tailed Student’s t-test with respective controls. D) Visual learning performance of adult flies with intact antennae upon silencing of visual KC subtypes (left: γd-KC, light blue, right: αβp-KC, dark blue) using Kir2.1. E) Same as (D) but in adult flies with their antennae cut. “n.s.”p>0.05, *p<0.05, ***p<0.001, two-tailed Student’s t-test. Boxplots in (C, D, E) show median, interquartile ranges and whiskers extend to the most extreme data points, excluding the outliers. Sample sizes for (C) α′β′m: n=8, α′β′ap: n=17, αβc: n=12, αβs: n=16, γm: n=14, γd: n=16, αβp: n=15 (flies); for (D) γd, empty>>Kir2.1: n=19, γd-KC>>Kir2.1: n=18, αβp, empty>>Kir2.1: n=8, αβp-KC>>Kir2.1: n=9 (groups of flies); (E) γd, empty>>Kir2.1: n=16, γd-KC>>Kir2.1: n=16, αβp, empty>>Kir2.1: n=13, αβp-KC>>Kir2.1: n=13 (groups of flies).

To assess changes in activity across KC subtypes, we used the long-term activity reporter CaLexA (calcium-dependent nuclear import of LexA) (Masuyama et al., 2012), allowing us to quantify GFP signals in KCs as a proxy for cumulative activity across the seven days following antennal removal (Fig. 3B). Using subtype-specific drivers for olfactory KCs or visual KCs, we showed that sensory deprivation by ablation of the antennae led to a selective shift in KC activity (Fig. 3C). Activity in some olfactory KCs, αβ-KCs (c and s) or γm-KCs, were unchanged, while it was reduced in olfactory α′β′-KCs (ap and m). In contrast, both visual KC populations (γd and αβp) showed increased activity (Fig. 3C). These results demonstrate that olfactory deprivation enhances activity of KCs that normally receive visual input, but not those that receive olfactory input.

To test how these changes relate to visual learning, we functionally blocked specifically these KC subtypes by expressing the inward-rectifier potassium channel Kir2.1 (Baines et al., 2001). Consistent with earlier studies (Vogt et al., 2014), blocking γd-KCs in intact flies abolished visual learning (Fig.3D, left). However, blocking αβp-KCs had no effect on visual learning in intact flies (Fig. 3D, right). In sensory-deprived flies, blocking γd-KCs again impaired visual learning (Fig. 3E, left). Strikingly, blocking αβp-KCs, KCs that are dispensable for visual learning in intact flies, disrupted memory performance in sensory-deprived flies (Fig. 3E, right). Further, blocking olfactory α′β′-KCs that showed reduced ongoing activity after sensory deprivation, did not change visual learning (Fig. S3A). Together, these results show that olfactory deprivation enhances activity of visual KCs in the adult and recruits an otherwise dispensable subtype of visual KCs, the αβp-KCs, into the memory circuit responsible for colour learning, highlighting the capacity of the MB to reconfigure memory storage by engaging additional visual pathways in response to olfactory loss.

### Olfactory deprivation releases visual γd-KCs from cross-modal suppression to enhance their responses to visual stimuli

In intact flies as well as in olfactory-deprived flies, visual learning depends on γd-KCs. Previous studies have shown that γd-KCs exhibit excitatory responses to colours (Vogt et al., 2016). We therefore expressed a genetically encoded calcium indicator in γd-KCs and tested whether their responsiveness to visual or olfactory stimuli changes after olfactory deprivation (antennal ablation, Fig. 4A). Flies were exposed sequentially to a visual stimulus, odour stimuli and a combination of olfactory and visual stimuli. Imaging visual responses in axons of γd-KCs in the lobes of the MB showed comparable responses in sensory-deprived and intact flies (Fig. 4B-C). In line with previous results (Vogt et al., 2016), testing for odour responses in γd-KCs revealed a slight reduction of calcium transients in intact flies (Fig. S4A). As expected, this reduction is not observed in olfactory-deprived flies without antennae (Fig. S4B). In line with this cross-modal interaction, we found that, when the visual cue was given in the background of an odour stimulus in intact flies, visual responses were reduced (Fig. 4B). However, this modulation was absent in olfactory-deprived flies (Fig. 4C). These findings suggest that olfactory input suppresses visual responses in γd-KCs.

**Figure 4:**
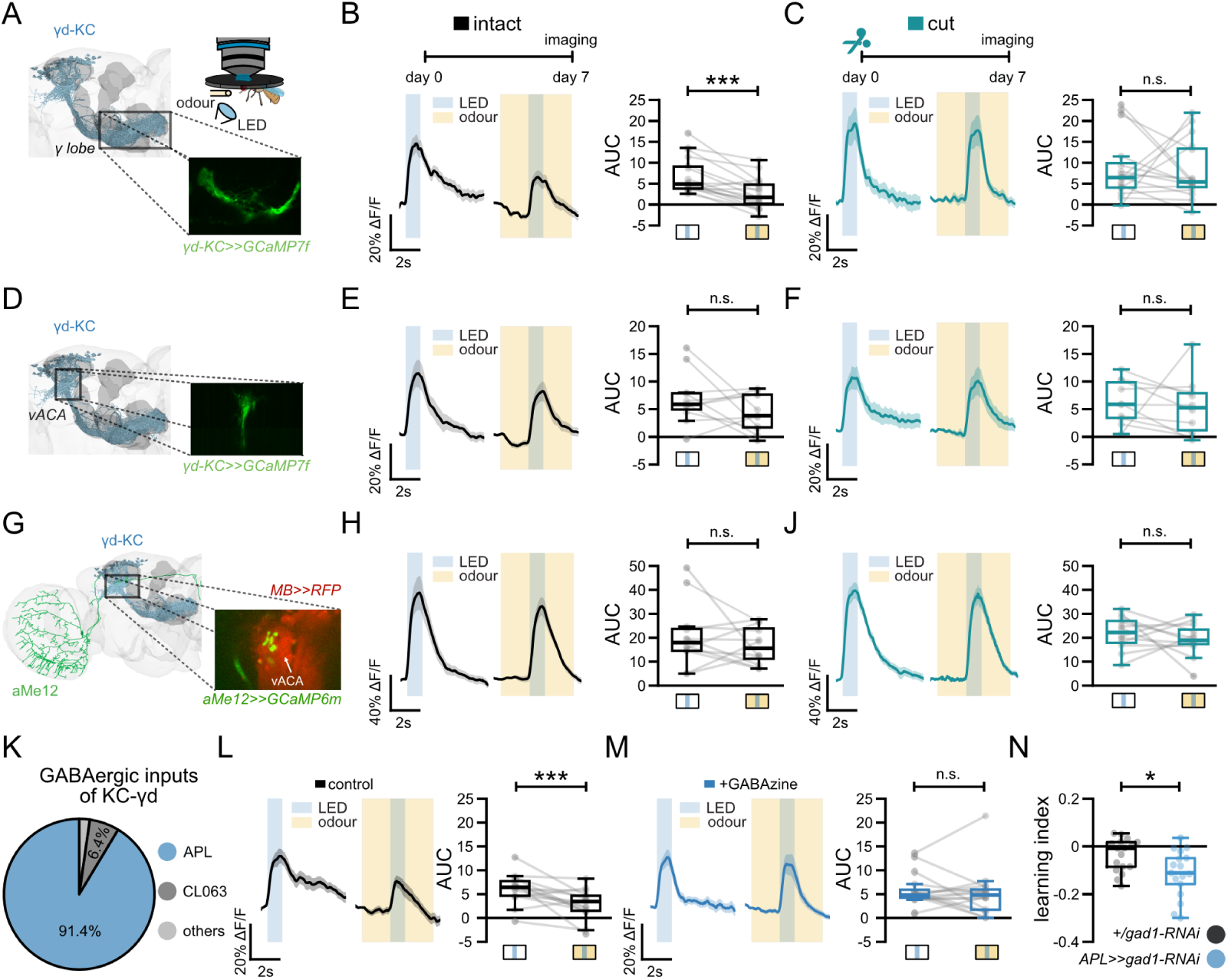
Olfactory deprivation releases γd-KCs from cross-modal suppression. A) Schematic overview of calcium imaging of visual and olfactory responses in axons of γd-KCs using the calcium indicator GCaMP7f. B) Calcium responses of γd-KCs in the MB lobes upon LED stimulation or LED stimulation in the background of an odour. Comparison of the response area under the curve (AUC) between the two stimuli. The flies have intact antennae. C) Same as (B) but in adult flies with their antennae cut. D, E, F) Same as (A, B, C) but calcium responses of γd-KCs are imaged at their dendrites at the ventral accessory calyx (vACA). G) Imaging of visual responses in γd-KCs’ input neuron aMe12-VPN at the vACA (identified by RFP expression using a MB driver line). H, J) Same as (B, C) but calcium responses of aMe12 neurons (green). K) The proportion of GABAergic inputs of γd-KCs based on the FAFB/FlyWire connectome. Shown neurons are the major inputs: APL (blue), CL063 (dark grey). L) Calcium responses of γd-KCs in the MB lobes before the application of GABA_A_ receptor inhibitor Gabazine with the quantification of AUC. M) Same as (L) but after the Gabazine application. N) Learning index after visual aversive conditioning of flies that express RNAi against gad1 to disrupt GABA production in APL neurons (blue) compared to RNAi control (black). Traces show mean ± SEM. “n.s.”p>0.05, *p<0.05, ***p<0.001, two-tailed Student’s t-test with paired (paired samples are connected via lines) or independent samples. Boxplots in (B, C, E, F, H, J, L, M, N) show median, interquartile ranges and whiskers extend to the most extreme data points, excluding the outliers. Sample sizes for (B, C) intact: n=16, cut: n=18 (flies); (E, F) intact: n=11, cut: n=10 (flies); (H, J) intact: n=12, cut: n=12 (flies); (L, M) control: n=15, gabazine: n=16 (flies); (N) +/gad1-RNAi: n=17, APL>>gad1-RNAi: n=17 (groups of flies).

Inhibitory interaction between KCs within the MB has been shown to be differentially regulated at distinct compartments of the neurons (Amin et al., 2020; Lei et al., 2013; Lin et al., 2014). To localize the source of inhibition, we imaged γd-KC dendrites in the calyx (Fig. 4D). We observed that the dendrites exhibited similar responses to the visual cue and a decay of calcium transients to odours (Fig. 4E, Fig. S4C-D). However, in contrast to the axonal responses in the lobes, dendrites of γd-KCs in the calyx showed no odour-driven suppression of visual responses (Fig. 4F). Furthermore, imaging the main input to γd-KCs, the aMe visual projection neurons (aMe-VPNs) (Ganguly et al., 2024), at the ventral accessory calyx (vACA) revealed excitatory visual responses but no odour-driven suppression in both intact and sensory-deprived flies (Fig. 4G-J). These results suggest that odour-evoked inhibition acts differently at the dendrites and the axons of the γd-KCs but not in upstream neurons, resulting in a robust olfactory suppression of visual responses in γd-KCs axons.

A candidate for such cross-modal inhibition is the GABAergic anterior paired lateral (APL) neuron, which mediates localized inhibition among KCs (Honegger et al., 2011; Lin et al., 2014; Masuda-Nakagawa et al., 2014). Consistent with this, assessing the FAFB/FlyWire connectome confirmed that GABAergic input to γd-KCs stems almost entirely from the APL neurons (Fig. 4K). To functionally test GABAergic inhibition, we recorded activity of γd-KCs axons in the absence and presence of the GABA*_A_* receptor antagonist Gabazine. While the presence of Gabazine did not substantially change visual responses in axons of γd-KCs (Fig. 4L-M), the presence of the antagonist abolished the odour-driven inhibition of visual responses (Fig. 4M), supporting the idea of GABAergic cross-modal inhibition. If this GABAergic inhibition restricts visual learning, then impairing this pathway should mimic the effect of sensory deprivation. To test this, we expressed RNAi against glutamic acid decarboxylase 1 gene (*gad1*) in APL neurons (VT243924-GAL4) (Amin et al., 2020; Dietzl et al., 2007). Targeting GABA synthesis in APL neurons does not change behaviour in mock conditions (Fig. S4E) but significantly enhanced visual learning in trained flies (Fig. 4N), phenocopying the effect of antennal ablation (Fig. 1G).

Together, these findings show that olfactory input suppresses visual responses in γd-KCs, apparently through localized GABAergic inhibition in the MB lobes. The GABAergic inhibition is most likely mediated by the APL neuron and restricts visual memory capacity. Olfactory deprivation reduces this axonal cross-modal inhibition and therefore enhances visual memory acquisition.

### Olfactory deprivation modulates the visual stream to αβp-KCs to enhance their responses to visual stimuli

Olfactory deprivation recruits αβp-KCs into visual memory circuits (Fig 3E). However, it remained unclear whether αβp-KCs respond directly to visual input and if these responses are changed after sensory deprivation. To address this, we imaged calcium responses of αβp-KCs’ dendrites in the dACA to a visual cue (Fig. 5A). In intact flies, αβp-KCs exhibited only marginal responses to visual cues (Fig. 5B). By contrast, in olfactory-deprived flies, these responses were enhanced, indicating that olfactory deprivation increases the visual input to αβp-KCs (Fig. 5B-C). To test whether this increase arose from MB–intrinsic mechanisms or from enhanced input via upstream visual projection neurons (VPNs), we examined VPNs that provide input to the dendrites of αβp-KCs. Previous studies identified the SLP448 neurons (referred to here as loVPN) as a major presynaptic partner of αβp-KCs(Ganguly et al., 2024; Li et al., 2020). Imaging of loVPN (R72D07-GAL4, GCaMP6m, with RFP in KCs) in the dACA revealed robust responses to visual cues in intact flies which are enhanced in olfactory-deprived flies (Fig. 5D-F). These findings demonstrate that olfactory deprivation enhances visual input from loVPNs to αβp-KCs. Together, these observations suggest that, in addition to the within-MB effects on γd-KCs, olfactory deprivation of antennal senses induces functional modulation outside of the MB that enhances αβp-KC responses to visual cues..

**Figure 5:**
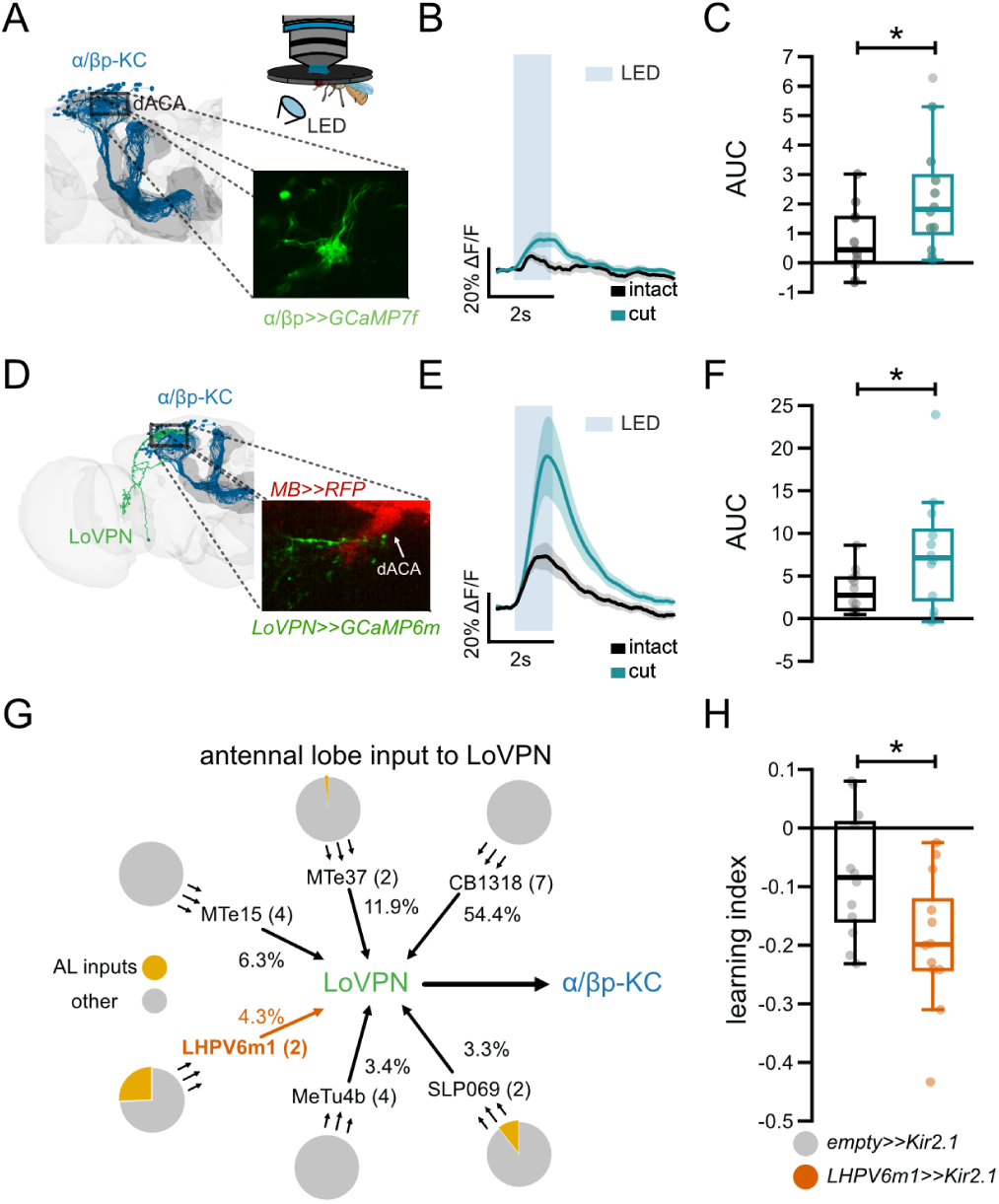
Sensory deprivation modulates the visual stream to αβp-KCs. A) Schematic overview of the imaging of visual responses of αβp-KCs in the dorsal accessory calyx (dACA). B) Calcium responses of αβp-KCs upon LED stimulation in flies with intact (black) and cut (teal) antennae. C) Comparison of AUC of αβp-KC calcium responses between intact (black) and cut (teal) flies. D) Schematic overview of the imaging of visual responses of LoVPNs in the calyx. E, F) Same as (B, C) but calcium responses of LoVPNs (green). G) Input connectivity map with percentage of inputs to LoVPN with their antennal lobe (AL) contribution (dark yellow) based on the FAFB EM dataset. The neuron with most AL inputs is LHPV6m1 (orange). H) Visual learning performance of adult flies upon silencing of the LoVPN input LHPV6m1 (orange) using cell-type specific expression of Kir2.1. Traces show mean ± SEM. *p<0.05, two-tailed Student’s t-test. Boxplots in (C, F, H) show median, interquartile ranges and whiskers extend to the most extreme data points, excluding the outliers. Sample sizes for B, C) intact: n=11, cut n=12 (flies); E, F) intact: n=12, cut n=12 (flies); H) empty>>Kir2.1: n=12, LHPV6m1>>Kir2.1: n=12 (groups of flies).

To identify the olfactory pathways that can intersect with loVPNs, we examined the antennal lobe inputs coming to loVPNs using the FAFB dataset. Amongst the presynaptic neurons of loVPNs, none have input synapses at the antennal lobe prompting us to investigate indirect pathways. From the top six input neurons of loVPNs, only the glutamatergic LHPV6m1 neuron gets substantial (∼26%) input from neurons that have input synapses at the antennal lobe (Fig. 5G). To test whether LHPV6m1 regulates visual learning, we expressed Kir2.1 (R64C05-GAL4) in the neurons. Blocking LHPV6m1 in intact animals increased visual learning, mimicking the effect of antennal removal (Fig. 5H), suggesting that pathways from the antennal lobe impinge on visual pathways to the MB, modulating visual learning.

Together, these findings suggest that olfactory deprivation recruits αβp-KCs to visual memory circuits through a changed control of the visual stream by antennal lobe input. This altered regulation seems to provide αβp-KCs with increased visual input, enabling their engagement in memory formation. The identification of the glutamatergic LHPV6m1 as a potential regulator of loVPNs will provide an entry point to map broader sensory interactions in healthy and deprived animals.

## Discussion

The brain is a highly plastic organ, capable of structural and functional remodelling in response to environmental changes, injury, or developmental disorders. A testament to this adaptability is cross-modal plasticity, through which animals adapt to the loss of one sensory modality by improving sensory processing of the remaining modalities (Merabet & Pascual-Leone, 2010). In some species, loss of one sensory modality has even been shown to improve higher cognitive functions, such as memory formation with intact sensory modalities (Amedi et al., 2003; Craig et al., 2022; Hori et al., 2006; Raz et al., 2007; Spigelman & Bryden, 1967). However, the synaptic and circuit mechanisms that underlie increased learning efficacy are not known. Here, we show that loss of olfactory sensory input enhances visual associative learning in larval and adult *Drosophila*. Using behavioural analysis, comparative connectomics, and imaging of neural activity, we found that improved learning is associated with stronger visual responses of neurons (Kenyon Cells [KCs]) in the higher-order learning circuit (the mushroom body [MB]). We discovered multiple, complementary circuit mechanisms that lead to enhanced responsiveness of KCs to visual stimuli: structural reweighting of inputs in the larva resulting in higher fraction of synaptic input from excitatory visual projection neurons (vPNs) or lower fraction of synaptic input from inhibitory visual local nuerons (vLNi) onto multisensory KCs; and reduced cross-modal inhibition in the adult at two distinct points in the circuit: onto visual γd-KCs and onto visual projection neurons (vPNs) presynaptic to visual αβp-KCs. Together, these changes reweight visual inputs to amplify visual representations in the memory centre and improve visual learning.

### Improvement of visual learning in the absence of olfaction

In humans, sensory loss can drive compensatory perceptual changes and, in some contexts, improve learning and memory in spared modalities. For example, congenital or early-onset blindness has been associated with enhanced performance in some auditory and verbal memory tasks (Amedi et al., 2003; Raz et al., 2007). Similarly, enucleated rats perform better in challenging auditory associative learning tasks than sighted controls (Spigelman & Bryden, 1967). By testing larval and adult *Drosophila*, we show that loss of olfactory input can enhance visual associative learning in an invertebrate. Notably, we demonstrate enhanced visual learning both under congenital deprivation (caused by mutation of the essential olfactory co-receptor gene *orco*) and following acute sensory organ damage later in life. These findings suggest that even compact brains can reallocate circuit resources after sensory loss, revealing mechanisms that support compensatory changes in learning.

### Increased Responsiveness of Learning Circuit Neurons to Visual Stimuli Enhances Visual Learning

How loss of one sensory modality could alter circuit structure and function in higher-order learning circuits to improve learning with remaining modalities is unknown. Our data support a convergent explanation: olfactory deprivation improves visual learning by increasing visual responses within the MB. In the larva, loss of olfaction enhances KC responses to visual stimuli (Fig. 2H). In adult flies, also, olfactory deprivation enhances visual responses in two visual KC populations. Although we discovered multiple different, complementary, circuit mechanisms that lead to these enhancements (structural reweighting of synaptic inputs resulting in increased input from excitatory visual vPNs and decreased input from inhibitory visual vLNi onto multisensory visual KCs in the larva; increased removal of cross-modal suppression to γd-KCs within the MB; and to vPNs presynaptic to αβp-KCs in the adult), all of these, ultimately, lead to increased responses of KCs that receive visual input to visual cues. Thus, across these circuit levels, olfactory deprivation converges functionally on increased sensitivity of KCs that receive visual input. Previous work has shown that increased KC responsiveness improves associative memory formation in the fly (Boto et al., 2019). Moreover, aversive visual learning, like olfactory learning, seems to depend on the coincidence of activity in punishment-induced dopamine neurons and activity of stimulus-encoding KCs (Vogt et al., 2014, 2016). Together, these findings suggest that increased activation of visual KCs ensures reliable coincidence detection, which in turn results in improved visual learning.

### Olfactory deprivation increases synaptic input from excitatory visual PNs onto multisensory neurons in the larval learning circuit

The *orco¹* mutant provides a genetic model of chronic olfactory deprivation, abolishing odorant receptor– dependent signalling (Benton, 2022; Fishilevich et al., 2005; Larsson et al., 2004). In the larval MB, two multisensory KCs receive visual input and integrate it with olfactory signals (Eichler et al., 2017). Through comparative connectomic analyses of two newly acquired EM volumes of larval brains (one *orco* mutant and the other wild-type control of the same age), we find that *orco* mutant larvae have increased fraction of synaptic input from a key excitatory vPN, vPN3, onto a specific KC, the *ov*KC, at the expense of olfactory inputs (Fig. 2H). This competitive reweighting of excitatory inputs contrasts with findings in the larval nerve cord, where inhibition of mechanosensory neurons resulted in increased connectivity from both the intact nociceptive pathway and from the deprived modality itself; a change consistent with homeostatic upregulation of excitatory inputs (Barnes et al., 2017; Bridi et al., 2018; Keck et al., 2017; Tripodi et al., 2008; Valdes-Aleman et al., 2021). Our findings in the higher-order learning circuit, suggest that the *ov*KC relies on competitive mechanisms between different excitatory inputs to determine connectivity, such that more active inputs are strengthened at the expense of inactive ones, in accordance with Hebbian rules (Abbott & Nelson, 2000; Brown et al., 1990; Malenka & Bear, 2004). In the larval nerve cord, increased fraction of excitatory synaptic input from sensory neurons correlates with increased responsiveness of downstream neurons to those sensory neurons (Valdes-Aleman et al., 2021). Similarly, the increase in fraction of visual excitatory input onto the ovKC observed here following olfactory deprivation is expected to enhance visual responsiveness of this KC. While we could not selectively image responses of the ovKC in the larva, imaging average responses of all KCs revealed increased responsiveness to visual stimuli in anosmic (orco mutant) larvae compared to controls (Fig 2H), consistent with the connectome.

Intriguingly, we did not observe an increase in synaptic connectivity from excitatory vPNs onto the *lon*KC, the other multisensory KC type that receives visual input in the larva. Unlike the ovKC, the lonKC receives not only excitatory, but also inhibitory visual input. Interestingly, we did observe a significant reduction in the fraction of synaptic input from an inhibitory visual local neuron (vLNi, Daniels et al., 2008; Larderet et al., 2017) onto the lonKC in anosmic animals. Thus, rather than increasing its excitatory input, *lon*KC may increase its responsiveness to visual stimuli by reducing feedforward inhibition from visual pathways.

### Reduction in Lateral Inhibition from Olfactory Pathways at Two Stages of Visual Processing Enhances Visual Responses of Learning Circuit Neurons in the Adult

Surgical removal of the antenna provides an acute, non-genetic perturbation that compromises multiple antennal modalities, including olfaction. Previous work indicates that antennal ablation can induce plasticity in the antennal lobe while preserving the gross organisation of projection neuron outputs(Berdnik et al., 2006; Kazama et al., 2011). Consistent with this, we find that overall MB anatomy and PN innervation patterns are preserved after antennal removal (data not shown), arguing against large-scale rewiring as a requirement for the behavioural change. Instead, antennal removal robustly enhances visual learning and visual KC responsiveness in the adult through two complementary circuit-level mechanisms: reduced cross-modal inhibition within the γ-lobe of the MB and increased engagement of αβp-KCs, which we trace to a change in upstream visual pathways that intersect with input from the antennal lobe.

Both γ KCs and αβ-p KCs receive visual input (Li et al., 2020; Vogt et al., 2016), but they differ in input pathways and default contribution to colour learning. The visual input to γd-KCs predominantly comes from direct visual projection neurons (Li et al., 2020). Based on this input pattern it has been suggested that they encode general features of the visual scene such as luminesces and colour and therefore provide a default substrate for associative learning in these settings (Ganguly et al., 2024; Li et al., 2020; Vogt et al., 2016). The odour-driven inhibition of this default visual learning pathway seems to be MB intrinsic. Effects of antennal ablation on γd-KCs are consistent with a loss of inhibitory control normally imposed by olfactory KCs, likely mediated via the APL neuron, which provides global and local inhibition across KC populations (Amin et al., 2020). In mice, the local removal of lateral inhibition between sensory pathways on the level of the thalamic reticular nucleus has been proposed as a possible mechanism to amplify signals to the cortex allowing heightened sensitivity within the remaining senses (Whitt et al., 2022). Thus, though it remains difficult to draw direct comparisons between brain structures in different animals, reducing local cross-modal inhibition at different functional levels appears to be a general mechanism for increasing the sensitivity of the remaining senses.

In contrast to γd-KCs, the functional role of adult αβp-KC is less well defined. Based on their input patterns αβp-KCs receive predominantly input from indirect visual pathways mediated by local visual interneurons, which convey highly processed and multisensory information from the optic lobes (Li et al., 2020). Therefore, it has been suggested that these neurons encode discrete objects and complex visual contexts (Li et al., 2020). Consistent with this idea, behavioural studies have shown that αβp-KC are involved in guiding visual attention toward dark objects such as stripes (Koenig et al., 2016). Together, these observations suggest that the enhancement of visual learning during sensory compensation is achieved through two complementary mechanisms: first, by strengthening the engagement of default visual learning pathways; and second, by recruiting functionally related multisensory pathways. This recruitment of functionally related pathways is reminiscent of compensatory mechanisms described after loss of vision in mammals (Petrus et al., 2014, 2015; Whitt et al., 2022). However, further work is required to understand how visual information is encoded across different KC subtypes and how these representations are changed during compensation.

In addition to odour detection, the antenna houses receptors for mechanosensation, thermosensation, hygrosensation, and other modalities, and plays a broad role in regulating physiology and behaviour (Dhar et al., 2024). Consequently, antennal removal compromises more than olfaction alone. Because αβp-KCs receive a larger fraction of their input through indirect and multisensory pathways, their increased visual responses could be influenced by the loss of other antennal senses beyond olfaction. Notably, hygrosensation and water-seeking behaviours critically depend on antennal sensory neurons and specifically engage αβp KCs (Ji & Zhu, 2015). We find that changes in αβp-KCs responses after sensory deprivation are inherited from modulation of visual pathways outside of the MB. Thus, disruption of antennal sense appears to rebalance multisensory inputs upstream of the MB, indirectly favouring visual information flow into αβp-KCs. Given the diversity of sensory modalities affected by antennal surgery, it is likely that additional compensatory mechanisms take place beyond the visual pathway to the MB. In line with this idea, recent work suggests that antennal removal alters gustatory processing and engages dopaminergic neurons innervating the MB, pointing to changes in reinforcement and modulatory systems as part of the adaptive response (Junca et al., 2021).

Together, our findings indicate that enhanced visual learning following antennal removal in the adult emerges from adjustments at multiple circuit levels: disinhibition within the MB and changes of visual pathways upstream of the MB. In line with contemporary frameworks of cross-modal plasticity, these results support a model in which compensation is dominated by functional (and structural) reweighting of existing circuits rather than large-scale anatomical rewiring that would generate new connections.

### Multisensory Neurons Enhance Responses to Spared Modality at the Expense of the Lost One

Recent work in vertebrates suggests that sensory enhancement after deprivation often involves changes in pre-existing multimodal circuits (Kral & Sharma, 2023; Lee & Whitt, 2015). Whether similar principles explain compensatory changes in cognitive tasks is less clear. Flies can learn from multisensory cues, such as combined visual and olfactory stimuli, and this multisensory learning, as shown in other animals (Shams & Seitz, 2008), outperforms unimodal learning (Thiagarajan et al., 2022). One possibility is that these gains arise through altered interactions between visual and olfactory KC populations and the recruitment of additional KC subtypes into the memory circuit. Our findings after sensory deprivation parallel this logic: in adults we see recruitment of αβp-KCs. In larvae our EM reconstructions show that a multisensory KC shifts strongly toward visual input at the expense of olfactory input in anosmic animals, consistent with activity-dependent Hebbian mechanisms that regulate the relative weighting of sensory modalities converging onto multisensory KCs.

Overall, our results suggest that compensatory changes in learning are implemented by modifying MB circuits that canonically support multisensory integration. This supports the broader view that cross-modal plasticity exploits existing connectivity between sensory pathways, and that understanding multisensory circuit structure is key to understanding adaptation after sensory deprivation.

### Broader Implications for Evolution of Improved Learning Capacity

Ecological demands shape both peripheral sensory systems and central learning circuits. In *Drosophila melanogaster*, olfaction is thought to be of crucial ecological importance for behaviours such as foraging and egg laying (Keesey et al., 2019; Mansourian et al., 2018). Previous studies indicate that associative olfactory learning is more robust than visual associative learning (Schnaitmann et al., 2013; Vogt et al., 2014). This behavioural asymmetry mirrors MB anatomy: most KCs receive olfactory input, whereas only a minority are driven by visual pathways (Eichler et al., 2017; Li et al., 2020). Our findings show that when olfactory input is compromised, visual circuits can assume a stronger role in associative learning. This supports a flexible hierarchy among modalities within the MB, in which olfaction typically dominates, but priority can be reassigned when inputs change. Our findings also reveal several circuit mechanisms that can lead to significant improvements in learning.

At an evolutionary scale, vision-olfaction trade-offs are also apparent: across *Drosophila* species, neural resource allocation in visual versus olfactory systems varies inversely, suggesting repeated, independent shifts in sensory prioritisation (Keesey et al., 2019). It is therefore tempting to speculate that the circuit mechanisms that improve learning after olfactory loss reflects circuit mechanisms that, under sustained ecological change, could be subject to selection and contribute to evolutionary shifts in learning ability.

## Methods

### Fly husbandry

All flies were reared on a cornmeal-based diet and maintained under controlled conditions of 25°C, 60% relative humidity and a 12:12h light:dark cycle. Parental fly stocks were reared at 20-21°C. Immunohistochemistry experiments were performed on 3-7d-old female flies. Physiological experiments (live imaging, CaLexA assays, and structural quantifications) were conducted on 7-day-old (±1) female flies unless otherwise stated. Behavioural experiments were performed on mixed-sex groups of 7-day-old (±1) flies. Larval experiments utilized *Canton-S* and *orco¹* strains reared on molasses food supplemented with retinal at 18°C; they were tested at either the early L2 (3–4 days old) or L3 feeding stage (7 days old). All fly lines are detailed in the **Key Resources Table**.

### Sensory deprivation in adult flies

Within 24 hours of eclosion, flies were anesthetized with CO_2_. For the experimental group, both antennae were completely removed using sterilized fine forceps/scissors, while the control group remained intact. Flies recovered for seven (±1) days under standard conditions.

### Behavioural experiments

#### Larval visual learning assay

Larval behavioural experiments used a behavioural setup described in *Croteau-Chonka et al.* with minor modifications (Croteau-Chonka et al., 2022). In brief, freely moving larvae were placed on 23 x 23 cm, 1.5 % agarose plates and presented with repeated sets of visual stimuli delivered by a digital micromirror device (520 nm, 100 uWcm-2) and punishing heat stimuli delivered via a targeted infrared laser (32.5oC, 1470nm). 60 s before training began, all larvae were exposed to three 20 s light exposures to acclimatise to the visual stimulus. To isolate the associative effects of training from habituation, two training paradigms were used. In the forward-paired paradigm, larvae were exposed to a 30 s light stimulus, with heat punishment delivered in the final ≈14 s. The backward-paired paradigm corresponded to ≈14 s of heat punishment terminating 5s before the onset of a 30 s light stimulus. These sets of pairing were repeated 20 times, with 105 s between the onset of stimuli in consecutive trials. 120 s after the offset of the final trial, larvae were exposed to three 20 s light exposures, separated by 45 s gaps where no stimulus was delivered. Larvae were tracked during this testing-phase using an adapted version of the Multi-Worm Tracker software (Ohyama et al., 2013), and a continuous read out of their behavioural choices was obtained using an incremental supervised learning model based on human tagging of larval videos (Masson et al., 2020).

Turning rates for each individual larva were extracted frame-by-frame and binned into 1 s windows. Larval identity throughout the three testing trials was resolved manually using a custom web-based interface. Only larvae that were tracked for at least 80% of all three testing trial periods (from 10 s before the testing stimulus onset to 10 s after the testing stimulus offset) were included in our analysis. Larvae respond to aversive and appetitive stimuli by altering their turning rates at the onset and offset of exposure (Croteau-Chonka et al., 2022; Gepner et al., 2015; Gomez-Marin et al., 2011; Klein et al., 2017). Therefore, we extracted a behavioural preference index (BPI) for each larva as (T_off_-T_on_)/2, where Ton and Toff are the mean change in turning rates at the stimulus onset (from the 10 s before stimulus onset to the 10 s after stimulus onset) and the mean turning change at stimulus offset (from the 10 s before stimulus offset to the 10 s after stimulus offset), respectively. This index gives positive values for approach-like turning behaviours and negative values for avoidance-like turning responses. By comparing the BPI values between forward-paired and backward-paired groups, we calculated a behavioural learning index (BLI) using the formula (BPI_forward_ - BPI_backward_)/2. Positive values for BLI reflect appetitive learning and negative values reflect aversive learning.

#### Larval odour preference assay

The olfactory preference of *orco* mutant and wild-type larvae were tested using an adapted version of the previously published Y-maze setup, where CO2 was replaced by natural odours (Lesar et al., 2021). Larvae were raised for 3 days at 25oC (late L2-stage), then placed in the Y-maze chamber. When the larva reaches the centre of the maze, it is given a choice between an arm containing an odour or the other one containing only air. They were each given 10 choices between 1-butanol (concentration of 10-2) and air. Their choices were recorded using an overhead camera and quantified using custom Python3 software.

#### Adult odour preference assay

Groups of 30–50 7-day-old flies were introduced into a custom-built T-maze. Using an elevator mechanism, flies were guided to a choice point between mineral oil and a 1:1000 dilution of 4-methylcyclohexanol (MCH). The assay was conducted in complete darkness with a 1 mL/s airflow per arm. The preference index (PI) was calculated as (Nodour − Ncontrol) / (Ntotal).

#### Adult visual learning assay

Visual conditioning was performed in custom arenas (Fig. 1A) equipped with transparent shock grids and cameras (FLIR Flea®3 FL3-U3-13E4C-C) for simultaneous stimulus delivery and video monitoring. The bottom of the arenas are made of glass with platinum wire electrodes with thicker quadrant wires (1 mm) and thinner inner quadrant wires (0.3 mm) separated by 1 mm. Visual stimuli were presented on an LCD monitor (5” HDMI LCD V2, Joy-It) positioned beneath the arenas, displaying blue (HEX #1D8AE1) and green (HEX #2D9932). Electric shocks were provided in 120 V DC and 10 mA for 1.5 s with an ITI of 3.5 s using a custom-built programmable stimulator synchronized by Stytra software (Štih et al., 2019).

Groups of 20–40 7-day-old flies were loaded into four parallel arenas. Reciprocal conditioning was performed simultaneously: one group received green paired with shock (CS+), while the other received blue (CS+). Training consisted of four alternating 1-minute CS+ and CS− presentations interleaved by 12 s of darkness. Following a 1-minute dark period, a shock was delivered 5 s before the test to promote fly behaviour. Preference was assessed by allowing flies to choose between colours for 1 minute (test phase 1), then flipping quadrant positions for another minute (test phase 2) to control for site bias. The final learning index (LI) was the average PI of the two reciprocal trials.

### Connectomics and connectome data processing

#### Larval connectomic analysis of visual inputs to the Mushroom Body

To collect a high-resolution volume of an *orco¹* and a wild-type larval brain, L2-stage larvae (three days at 18°C) were dissected and prepared based on fixation and staining protocols previously described, but with the adaptations detailed in the table below (Lu et al., 2022). They were imaged using an enhanced focused ion-beam scanning electron microscope (eFIB-SEM, landing energy 1.2 kV, beam current 3 nA, FIB voltage 30 kV, 30 nA milling probe – 14nA used for milling) (Xu et al., 2017), resulting in an isotropic resolution of 8 x 8 x 8 nm. The datasets were automatically segmented using a flood-filling network model described in Januszewski et al. (Januszewski et al., 2018). Manual proofreading and synapse tagging was performed on skeletonised versions of these segmentations using CATMAID (Saalfeld et al., 2009). The identification of neurons from proofread skeletons was achieved through comparisons to the larval connectome based on shared morphology as determined using NBLAST (Costa et al., 2016; Winding et al., 2023). Data analysis was performed using a combination of native CATMAID tools and custom Python scripts. We defined any neuron that provides greater than 3% input to a KC in both hemispheres of either the *orco^1^* or wild-type volume, as a ‘strong connector’. If a neuron was identified as a strong connector in only one of the volumes, we searched through the weak connectors (<3% input) in the other volume to identify it.

Staining protocol:

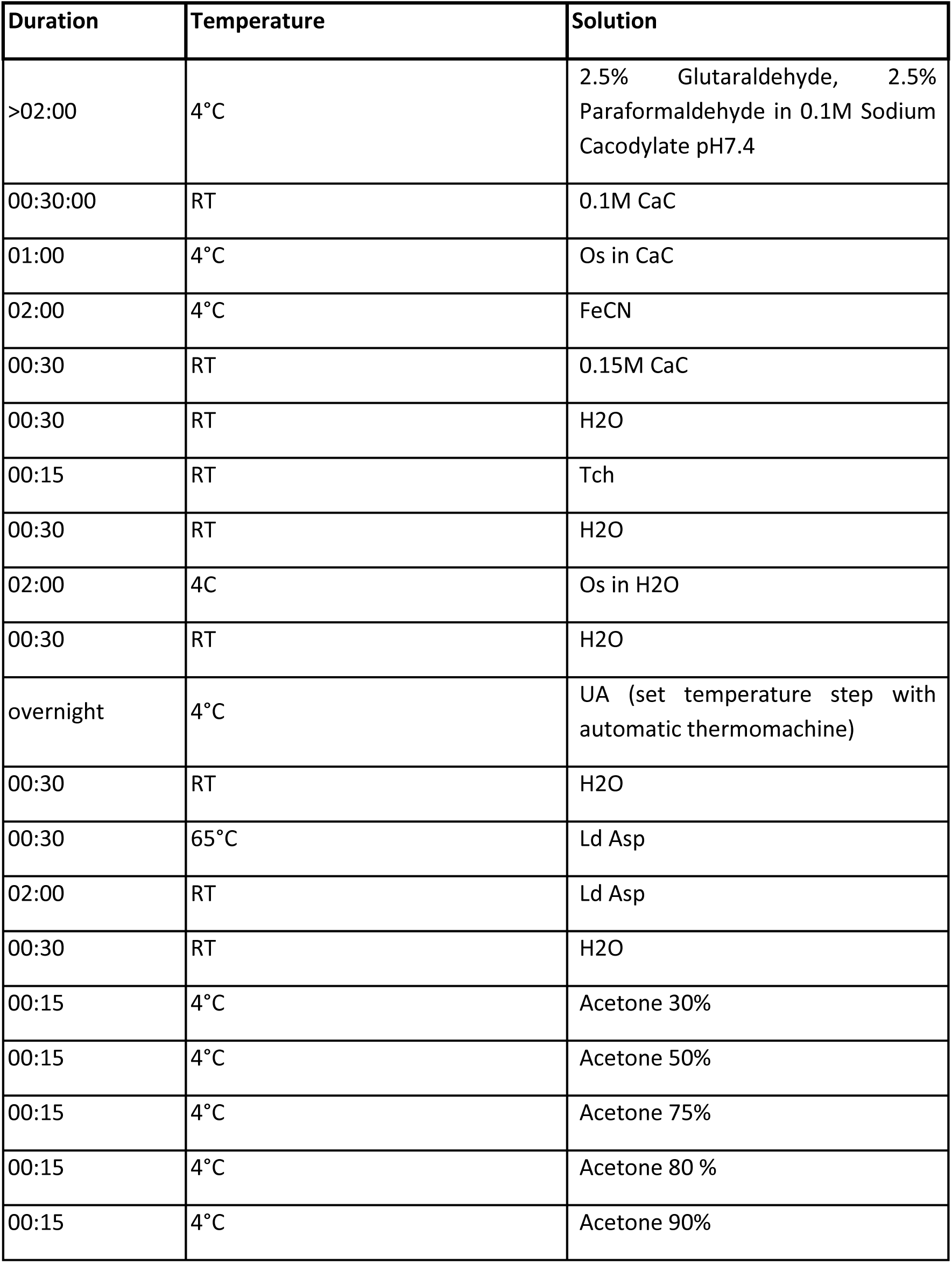
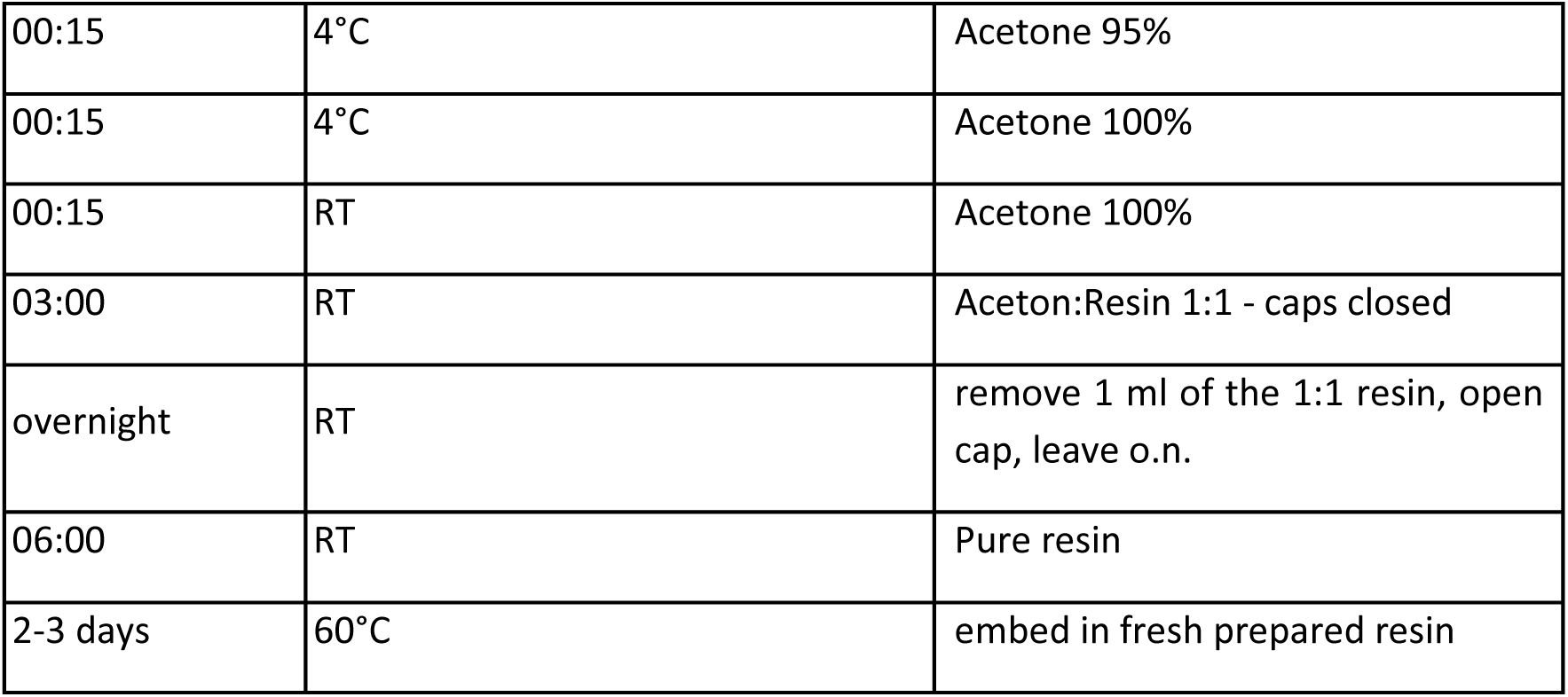

| Duration | Temperature | Solution |
| --- | --- | --- |
| >02:00 | 4°C | 2.5% Glutaraldehyde, 2.5% Paraformaldehyde in 0.1M Sodium Cacodylate pH7.4 |
| 00:30:00 | RT | 0.1M CaC |
| 01:00 | 4°C | Os in CaC |
| 02:00 | 4°C | FeCN |
| 00:30 | RT | 0.15M CaC |
| 00:30 | RT | H2O |
| 00:15 | RT | Tch |
| 00:30 | RT | H2O |
| 02:00 | 4C | Os in H2O |
| 00:30 | RT | H2O |
| overnight | 4°C | UA (set temperature step with automatic thermomachine) |
| 00:30 | RT | H2O |
| 00:30 | 65°C | Ld Asp |
| 02:00 | RT | Ld Asp |
| 00:30 | RT | H2O |
| 00:15 | 4°C | Acetone 30% |
| 00:15 | 4°C | Acetone 50% |
| 00:15 | 4°C | Acetone 75% |
| 00:15 | 4°C | Acetone 80 % |
| 00:15 | 4°C | Acetone 90% |
| 00:15 | 4°C | Acetone 95% |
| 00:15 | 4°C | Acetone 100% |
| 00:15 | RT | Acetone 100% |
| 03:00 | RT | Aceton:Resin 1:1 - caps closed |
| overnight | RT | remove 1 ml of the 1:1 resin, open cap, leave o.n. |
| 06:00 | RT | Pure resin |
| 2-3 days | 60°C | embed in fresh prepared resin |

Key:

|  |  |
| --- | --- |
| Os in CaC | 4% OsO<sub>4</sub> in h<sub>2</sub>O 1:1 volume ratio mix with 0.2 M CaC pH 7.4 |
| FeCN | 2.5% potassium ferrocyanide (Sigma) in 0.1 M CaC |
| Tch | 1% in h<sub>2</sub>O |
| Os in h<sub>2</sub>O | 4% OsO<sub>4</sub> in h<sub>2</sub>O 1:1 volume ratio mix with h<sub>2</sub>O |
| UA | 2% uranium acetate in h<sub>2</sub>O |
| Ld asp | 0.066 g Lead nitrate in 10 ml 0.03 M aspartic acid, pH adjusted to 5.0 with KOH |
| resin | Durcupan |

#### Adult connectome FAFB data analysis

Adult connectivity analysis was done using the FlyWire instance (v783) of the full adult female fly brain (FAFB) dataset via custom-written code in Python with the fafbseg Python package (Dorkenwald et al., 2024; Schlegel et al., 2024). Distribution of KC (variable cell_class: “Kenyon_Cell”) subtypes (variable cell_type) were done based on cell counts. For the GABAergic input analysis of KC-γd (295 KCs in FAFB/FlyWire), pre-synaptic partners with a minimum of 3 synapses were used. Determining the neurotransmitter profile was done using the top neurotransmitter prediction of the corresponding input neuron. Average synapses per KC were calculated before calculating the percentage of inputs. For the analysis of inputs to LoVPN (LTe72), cumulative synaptic counts of pre-synaptic partners were aggregated by partner type to first determine the percentage of synaptic contributions to LoVPN. From the inputs of these pre-synaptic partners, the contribution from the Antennal Lobe was determined via the fraction of synapses coming from neurons that have input synapses in the Antennal Lobe.

### Immunohistochemistry and confocal microscopy

#### Sample preparation and imaging

7d-old female flies (unless otherwise stated) were anesthetized on ice and were dissected in and transferred to chilled Ringer′s solution (130 mM NaCl, 5 mM KCl, 2 mM MgCl₂, 2 mM CaCl₂, 36 mM sucrose, 5 mM HEPES, pH 7.3). After fixation in 4% paraformaldehyde (PFA) for 1 hour, brains were washed three times (each lasting at least 15 minutes) in PBS containing 0.6% Triton-X-100 (PBS-T; 137 mM NaCl, 10 mM Na₂HPO₄, 2.7 mM KCl, 1.8 mM KH₂PO₄, pH 7.3), and incubated in blocking solution (2% bovine serum albumin in PBS-T) overnight at 4°C. Primary antibodies in blocking solution were applied for at least 4 hours, followed by PBS washes. Secondary antibodies were incubated overnight at 4°C, with subsequent washes. Brains were then mounted in VECTASHIELD, oriented as needed for imaging. Confocal images were acquired using a Yokogawa CSU W1 spinning disk unit on a Zeiss Axio Imager M2 microscope equipped with either a 20x air or a 40x oil immersion objective. A 488 nm laser (iBeam, Toptica Photonics, Germany) was used to excite GFP and a 561 nm laser (Juve, Hübner Photonics, Germany) was used to excite RFP. The setup was equipped with 460/50 and 525/50 emission filters. The images were obtained in 1 or 1.5 μm z steps. Image processing was performed using Fiji. Antibody details, sources, and dilutions are provided in the “Key Resources Table”.

#### Quantitative reporter expression analysis for CaLexA experiments

Seven-day-old (±1 d) female flies were used for experiments. Using Fiji, maximum projections of the calyx region were generated. Background fluorescence was measured from a selected region of interest (ROI) and subtracted from the entire stack to correct for background artifacts. Dendritic arborisations were isolated using thresholding, and the calyx ROI was defined. Autofluorescence regions above were excluded manually before measuring the fluorescence intensity. Data was analysed using GraphPad Prism 10.

### *In vivo* calcium imaging

#### Larval KC activity recordings

Volumetric fluorescence imaging of dissected larval Drosophila CNS preparations was performed using a SiMView light-sheet microscope as described previously (Lemon et al., 2015). Briefly, two opposing illumination arms (custom-made 10X, water dipping objectives, Special Optics) provided uniform excitation across the sample, while two opposing detection arms (20X, water dipping, Zeiss) acquired dual-view image volumes at ∼1 volume/s and 0.33 × 0.33 × 1 µm voxel resolution. Dual-view volumes were merged using custom Python software.

Visual stimulation was delivered using a 488 nm laser through one of the detection objectives, allowing spatially restricted stimulation of selected regions of the sample. Patterned stimulation was controlled by a Digital Micromirror Device-based optical path, allowing an in-plane resolution of ∼1 µm. Second instar larvae (raised for 4 days at 18°C) expressing the red-shifted calcium indicator, RGECO (Dana et al., 2016), in rh5 photoreceptors (Rh5-lexA) and Kenyon cells (14H06-lexA) were dissected in Baines’ solution, and the isolated CNS was embedded in low-melting-point agarose in a glass capillary. Samples were then transferred to the imaging chamber for fluorescence imaging. The imaging protocol consisted of an initial passive imaging phase: 561 nm light-sheet imaging was performed without external stimulation for 10 minutes. Subsequently, we performed the stimulation phase, in which five 10 s stimuli (488nm laser, average power ∼1mW/mm^2 measured at the objective back aperture) were delivered to the Rh5 target region at 60 s intervals, followed by five 20 s stimuli delivered under the same conditions. The mean response across the 20s stimulations were used for our analysis. The lobes of the MB were manually segmented using Napari (https://napari.org/).

#### Adult calcium imaging

Flies were anesthetized and mounted onto a custom platform, with the head capsule aligned to a viewing window and immobilized with hair strings across the neck and proboscis. The abdomen and head were secured with paraffin wax, a drop of hemolymph solution (5 mM TES, 103 mM NaCl, 3 mM KCl, 1.5 mM CaCl₂, 4 mM MgCl₂, 26 mM NaHCO₃, 1 mM NaH₂PO₄, 8 mM trehalose, 10 mM glucose; pH∼7) was applied to the head capsule, and the cuticle was opened to remove trachea and fat tissue. Samples were imaged on an upright microscope Axio Imager M2 (Zeiss) equipped with CSU-W1 Confocal Scanner Unit (Yokogawa) with a 50 μm pinhole disk unit, a sCMOS camera (PCO.edge 4.2M, PCO), an MS-2000 motorized stage (Applied Scientific Instrumentation) and controlled with VisiView imaging software (Visitron Systems GmbH). Illumination was achieved with 488 iBeam (Toptica Photonics) and 561 Jive (Cobolt) lasers and VS-Homogenizer (Visitron Systems GmbH). Plan Apochromat 40X/1.0 water immersion objective (Zeiss) was used. Emission filters in the scanhead were BP525/50 and BP609/54. Recordings were obtained using a laser power of approx. 7 mW for 488 nm laser and approx. 9mW for 561 nm laser and frames were acquired at 25 Hz.

Visual stimulation was done using an LED (Catalog#: 244 1400, CoolLED, England). LED-only trials had three repetitions of 1-s blue pulses (400 nm, 50% power) at fixed intervals. Olfactory stimulus was delivered directly to the antennae of the fly using an olfactometer (220A Aurora Scientific, Canada) at 900 mL/s airflow. Odour-only trials consisted of 3 s of odour presentation (3-Oct or MCH). Combined stimulation consisted of a 1-s LED pulse presented midway through a 5s odour pulse.

For the GABA_A_ blocking pharmacology experiments, Gabazine was prepared as a 10 mM stock solution in distilled water and stored at −20°C. On the day of imaging it was diluted to 50 µM in hemolymph solution. Each brain was first imaged in hemolymph solution without Gabazine and subsequently superfused with Gabazine-containing hemolymph to allow direct within-sample comparison of neuronal responses before and after GABA_A_ receptor blockade.

Time series of recordings were first motion corrected using the *template matching* plugin in Fiji followed by smoothing in space (2x) and time (3x) for noise reduction (Tseng et al., 2011). ROIs and the background regions were manually selected on the maximum intensity projection images. Following background subtraction, ΔF/F0 was calculated using the mean signal of a 1 s period before any stimulation occurred. After this, the analysis continued in custom-written Python code. The 3 trials of LED-only traces were averaged and stimulus specific analyses were performed for each fly. To calculate the area-under-the-curve (AUC), the calcium signals were integrated during the stimulation period using the composite trapezoid rule (scipy.integrate.trapezoid). Statistical tests are reported in each figure.

### Sleep profile analysis

Female flies were individually placed in 65 mm x 5 mm capillary glass tubes containing fly culture food and loaded into the Trikinetics *Drosophila* Activity Monitor system (DAM2). The activity monitors were maintained under 12h:12h light:dark conditions at 25°C and 65% relative humidity. Following a one-day habituation period, flies’ locomotor activity was recorded for five consecutive days. Activity data was collected in 1 minute bins, and periods of immobility lasting at least 5 min were classified as sleep (Hendricks et al., 2000; Shaw et al., 2000). Sleep data was analysed using the sleep and circadian analysis MATLAB program (SCAMP) (Vecsey et al., 2024). Sleep and activity parameters were quantified and averaged across four days of recording. Flies that were visibly dead or housed in tubes containing larvae were excluded from the analysis. Up to 16 flies per group were included in each experiment, and three independent experiments were done using flies from separate batches.

### Statistical analysis

Data were analysed in GraphPad Prism 10 or custom Python scripts. Each behavioural and imaging experiment was repeated at least three times to ensure reproducibility. Groups were compared using unpaired two-tailed Student’s t-tests, or paired t-tests for Gabazine experiments. Multiple comparisons were performed via one-way or two-way ANOVA with Tukey’s or Šidák’s post hoc tests.

## Key resource table

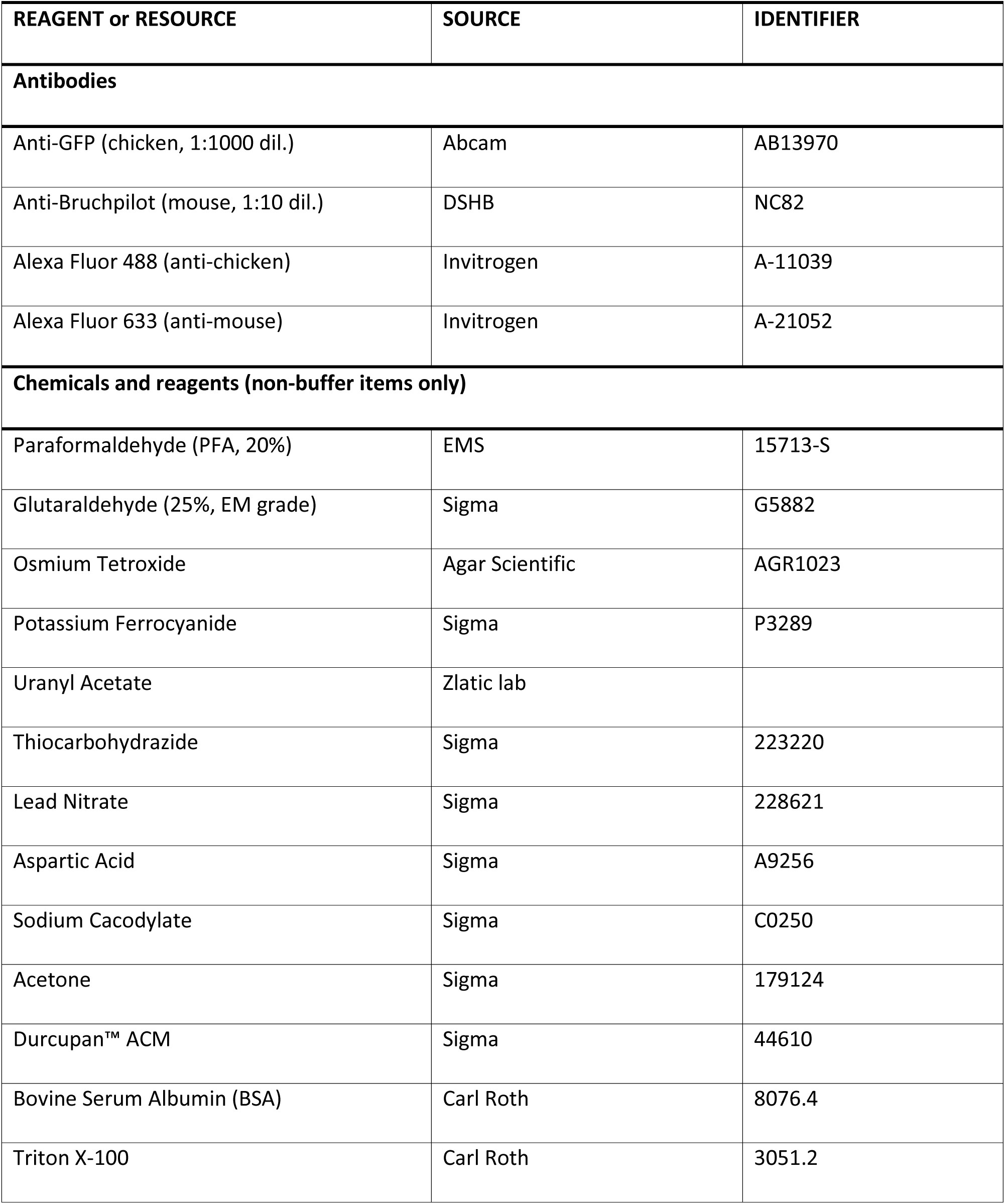
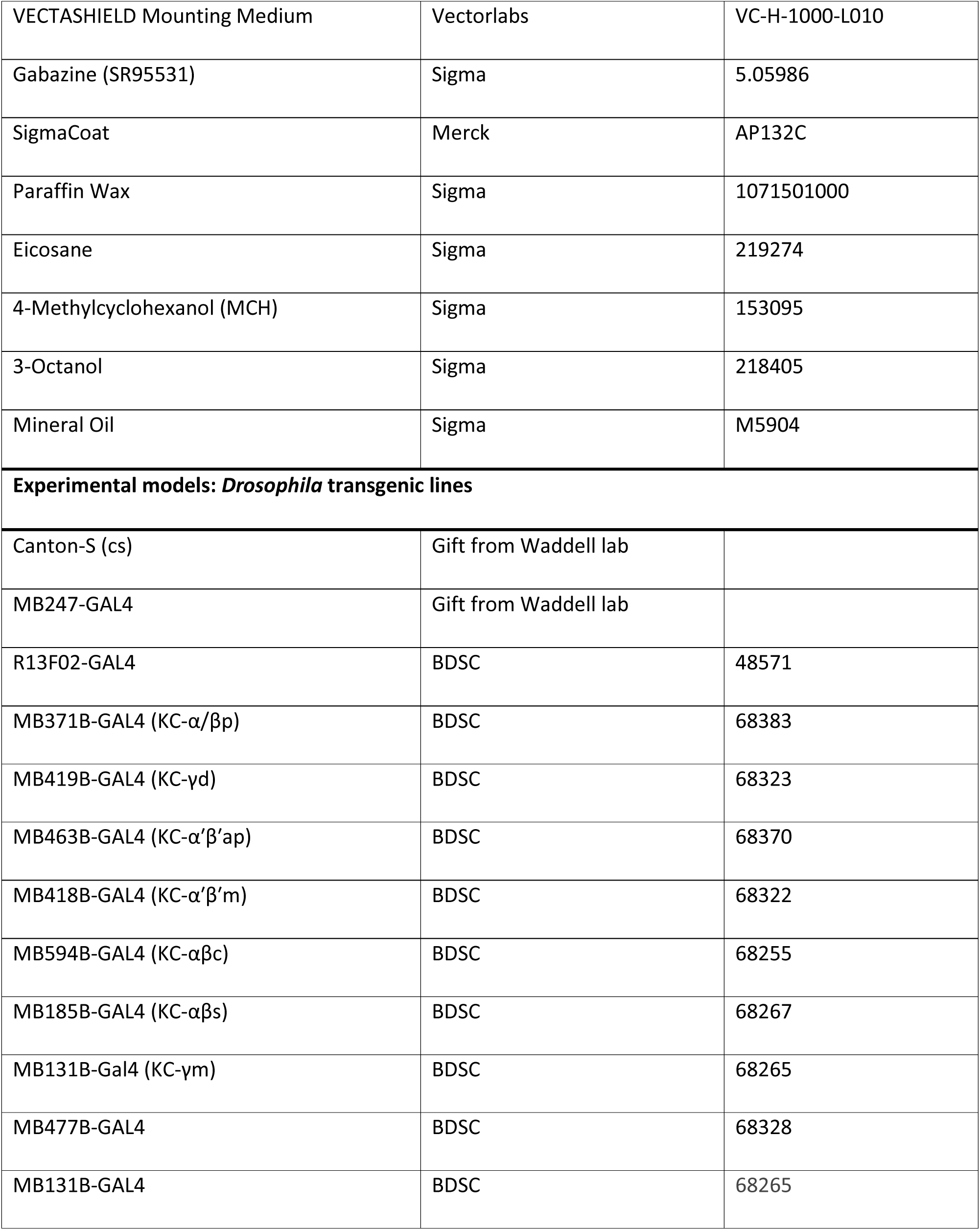
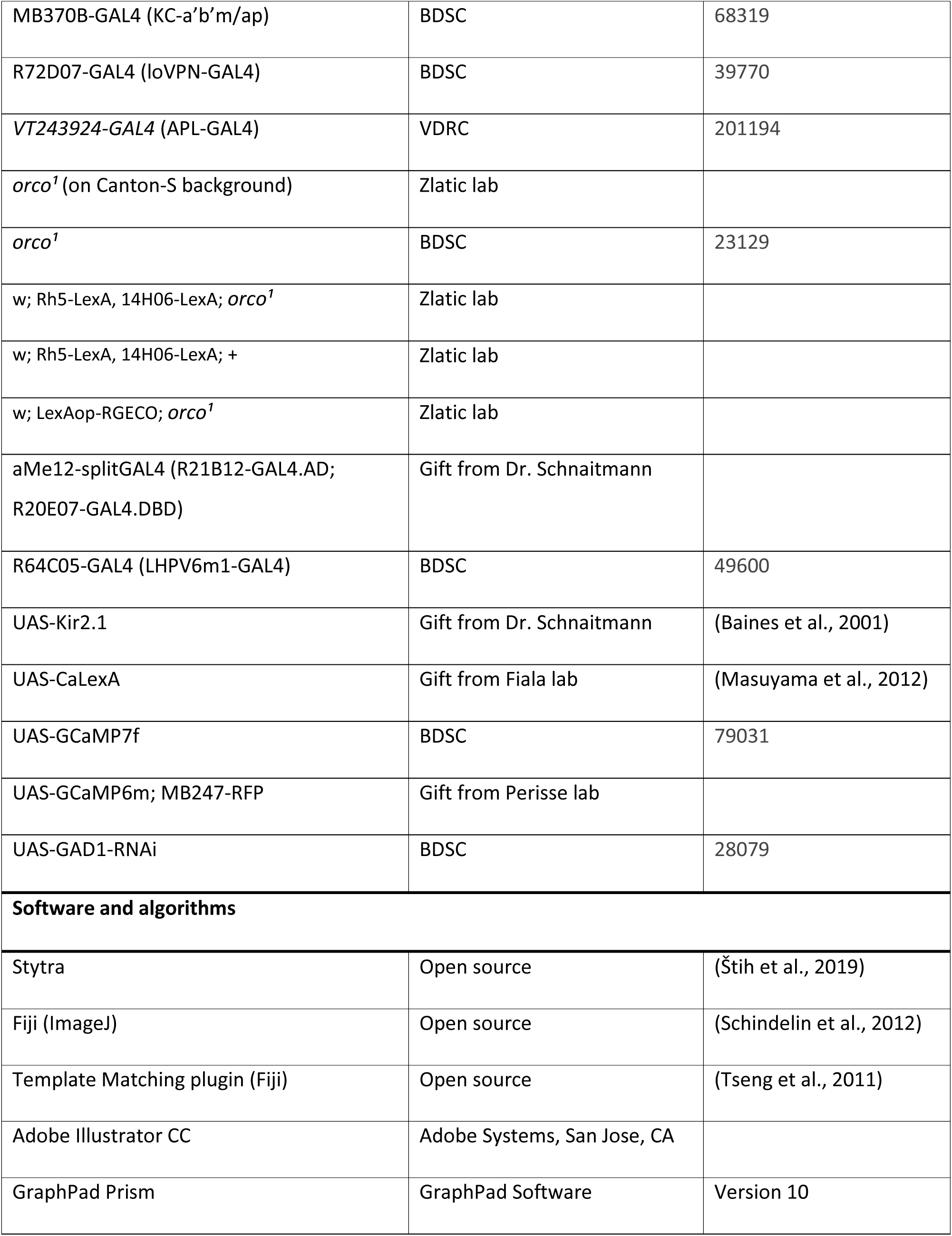
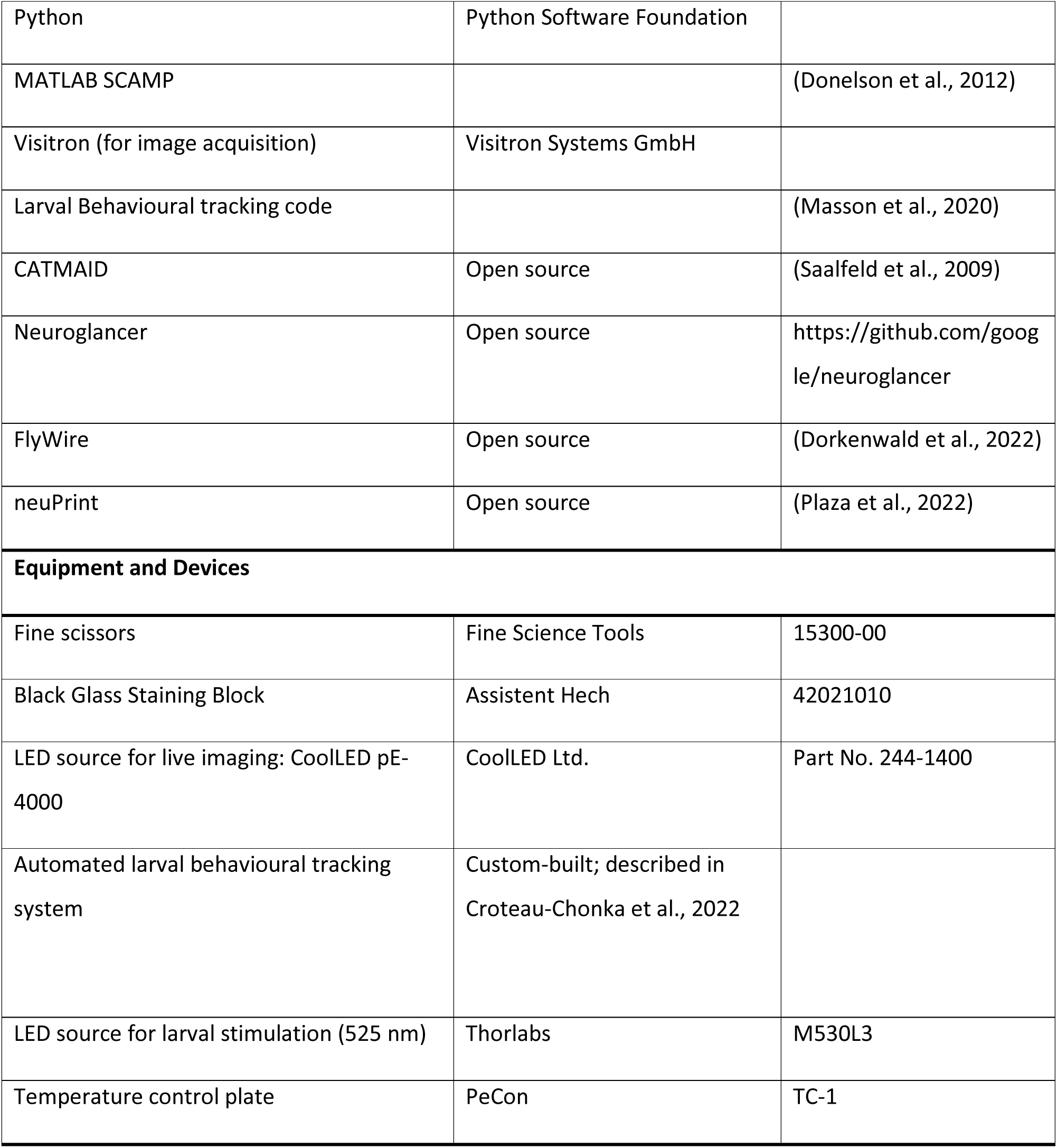

| REAGENT or RESOURCE | SOURCE | IDENTIFIER |
| --- | --- | --- |
| **Antibodies** |  |  |
| Anti-GFP (chicken, 1:1000 dil.) | Abcam | AB13970 |
| Anti-Bruchpilot (mouse, 1:10 dil.) | DSHB | NC82 |
| Alexa Fluor 488 (anti-chicken) | Invitrogen | A-11039 |
| Alexa Fluor 633 (anti-mouse) | Invitrogen | A-21052 |
| **Chemicals and reagents (non-buffer items only)** |  |  |
| Paraformaldehyde (PFA, 20%) | EMS | 15713-S |
| Glutaraldehyde (25%, EM grade) | Sigma | G5882 |
| Osmium Tetroxide | Agar Scientific | AGR1023 |
| Potassium Ferrocyanide | Sigma | P3289 |
| Uranyl Acetate | Zlatic lab |  |
| Thiocarbohydrazide | Sigma | 223220 |
| Lead Nitrate | Sigma | 228621 |
| Aspartic Acid | Sigma | A9256 |
| Sodium Cacodylate | Sigma | C0250 |
| Acetone | Sigma | 179124 |
| Durcupan™ ACM | Sigma | 44610 |
| Bovine Serum Albumin (BSA) | Carl Roth | 8076.4 |
| Triton X-100 | Carl Roth | 3051.2 |
| VECTASHIELD Mounting Medium | Vectorlabs | VC-H-1000-L010 |
| Gabazine (SR95531) | Sigma | 5.05986 |
| SigmaCoat | Merck | AP132C |
| Paraffin Wax | Sigma | 1071501000 |
| Eicosane | Sigma | 219274 |
| 4-Methylcyclohexanol (MCH) | Sigma | 153095 |
| 3-Octanol | Sigma | 218405 |
| Mineral Oil | Sigma | M5904 |
| **Experimental models: *Drosophila* transgenic lines** |  |  |
| Canton-S (cs) | Gift from Waddell lab |  |
| MB247-GAL4 | Gift from Waddell lab |  |
| R13F02-GAL4 | BDSC | 48571 |
| MB371B-GAL4 (KC-α/βp) | BDSC | 68383 |
| MB419B-GAL4 (KC-γd) | BDSC | 68323 |
| MB463B-GAL4 (KC-$\alpha'$ $\beta'$ap) | BDSC | 68370 |
| MB418B-GAL4 (KC-$\alpha'$ $\beta'$m) | BDSC | 68322 |
| MB594B-GAL4 (KC-$\alpha\beta$c) | BDSC | 68255 |
| MB185B-GAL4 (KC-$\alpha\beta$s) | BDSC | 68267 |
| MB131B-Gal4 (KC-γm) | BDSC | 68265 |
| MB477B-GAL4 | BDSC | 68328 |
| MB131B-GAL4 | BDSC | 68265 |
| MB370B-GAL4 (KC-a'b'm/ap) | BDSC | 68319 |
| R72D07-GAL4 (IoVPN-GAL4) | BDSC | 39770 |
| VT243924-GAL4 (APL-GAL4) | VDRC | 201194 |
| *orco*<sup>1</sup> (on Canton-S background) | Zlatic lab |  |
| *orco*<sup>1</sup> | BDSC | 23129 |
| w; Rh5-LexA, 14H06-LexA; *orco*<sup>1</sup> | Zlatic lab |  |
| w; Rh5-LexA, 14H06-LexA; + | Zlatic lab |  |
| w; LexAop-RGECO; *orco*<sup>1</sup> | Zlatic lab |  |
| aMe12-splitGAL4 (R21B12-GAL4.AD;<br>R20E07-GAL4.DBD) | Gift from Dr. Schnaitmann |  |
| R64C05-GAL4 (LHPV6m1-GAL4) | BDSC | 49600 |
| UAS-Kir2.1 | Gift from Dr. Schnaitmann | (Baines et al., 2001) |
| UAS-CaLexA | Gift from Fiala lab | (Masuyama et al., 2012) |
| UAS-GCaMP7f | BDSC | 79031 |
| UAS-GCaMP6m; MB247-RFP | Gift from Perisse lab |  |
| UAS-GAD1-RNAi | BDSC | 28079 |
| **Software and algorithms** |  |  |
| Stytra | Open source | (Štíh et al., 2019) |
| Fiji (ImageJ) | Open source | (Schindelin et al., 2012) |
| Template Matching plugin (Fiji) | Open source | (Tseng et al., 2011) |
| Adobe Illustrator CC | Adobe Systems, San Jose, CA |  |
| GraphPad Prism | GraphPad Software | Version 10 |
| Python | Python Software Foundation |  |
| MATLAB SCAMP |  | (Donelson et al., 2012) |
| Visitron (for image acquisition) | Visitron Systems GmbH |  |
| Larval Behavioural tracking code |  | (Masson et al., 2020) |
| CATMAID | Open source | (Saalfeld et al., 2009) |
| Neuroglancer | Open source | https://github.com/google/neuroglancer |
| FlyWire | Open source | (Dorkenwald et al., 2022) |
| neuPrint | Open source | (Plaza et al., 2022) |
| **Equipment and Devices** |  |  |
| Fine scissors | Fine Science Tools | 15300-00 |
| Black Glass Staining Block | Assistent Hech | 42021010 |
| LED source for live imaging: CoolLED pE-4000 | CoolLED Ltd. | Part No. 244-1400 |
| Automated larval behavioural tracking system | Custom-built; described in Croteau-Chonka et al., 2022 |  |
| LED source for larval stimulation (525 nm) | Thorlabs | M530L3 |
| Temperature control plate | PeCon | TC-1 |

## Declaration of interests

The authors declare no competing financial interests.

## Acknowledgments

The authors thank Nan Hu and Oxana Elliott at the Department of Zoology, University of Cambridge for assistance with fly crosses. Work in the Schnaitmann laboratory was supported by the German Research Foundation (DFG; grant SCHN1479/1-2). Work in the Kempf laboratory was supported by the Swiss State Secretariat for Education, Research and Innovation (SERI; M822.00025, SERI-funded ERC Starting Grant to A.K.) and by funds from the Cantons of Basel-Stadt and Basel-Landschaft. Work in the Zlatic laboratory was supported by core funding from the MRC Laboratory of Molecular Biology (M.Z., A.C., S.H., N.C., and M.C.), Wellcome Trust grant 205038/Z/16/Z (A.C., M.C., and A.J.C.S.), Wellcome Trust grant 205050/Z/16/Z (M.Z., I.H., and N.H.), European Research Council Consolidator Grant ERC-2018-COG-819650 (M.Z., A.D., and O.E.), and the LMB/AZ Blue Sky Fund Award BSF2-21 (K.W.). Work in the Felsenberg laboratory was supported by the European Research Council (ERC Starting Grant No. 950480) and the Swiss National Science Foundation (SNSF). B.G. was supported by a Human Frontier Science Program (HFSP) Postdoctoral Fellowship and D.G. by DFG Walter Benjamin Postdoctoral Fellowship. Stocks obtained from the Bloomington Drosophila Stock Center (NIH P40OD018537) were used in this study.

## Author contributions

**Conceptualization** BC, SNH, MZ, JF, **Methodology** BC, SNH, MZ, CS, AK, JF, AC, MC, AJCS, AC, MJ, AD, **Software** BG, SNH, DG, BC, AC, MJ, **Validation:** SNH, BC, BG, KL, SYL, **Formal analysis:** BC, SNH, BG, DG, DS, AK, **Investigation** BC, SNH, KL, DS, MC, AJCS, AC, IH, NC, **Resources** AC, MC, AJCS, MJ, **Data Curation** BC, SNH, BG, DG, CS, DS, SYL, AJCS, **Writing - Original Draft** SNH, BC, MZ, JF, **Writing - Review & Editing** BG, DG, SNH, BC, MZ, JF, **Visualization** SNH, BC, DG, BG, **Supervision** MZ, JF, **Project administration** BC, SNH, MZ, JF, **Funding acquisition** MZ, JF

---

**Figure S1:**
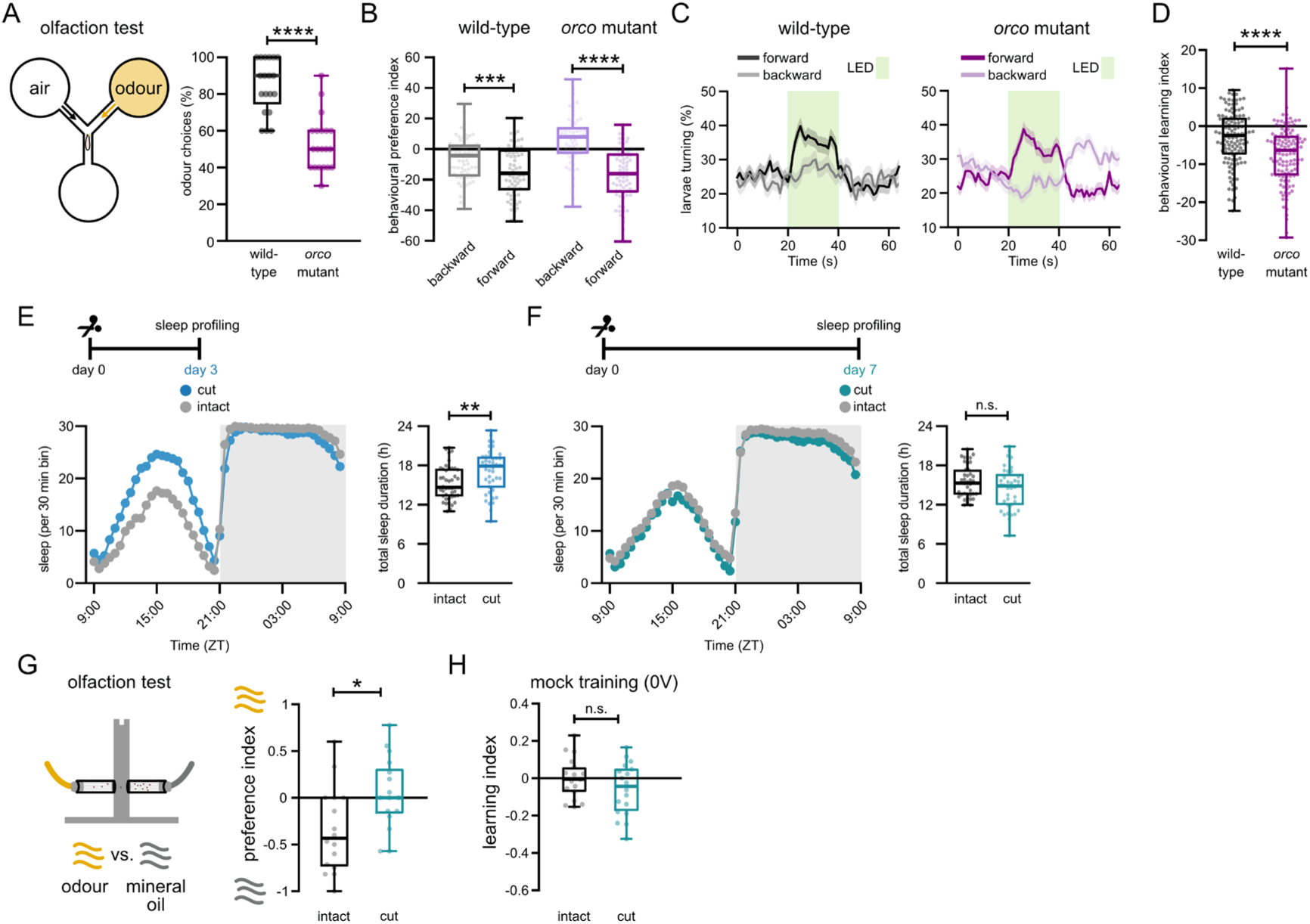
A) Left: Odour attraction assay to probe for olfaction in the larvae. Right: Comparison of the percentage of larvae choosing the odour side between wild-type (grey) and orco mutants (magenta). ****p<0.0001, Mann-Whitney U test. B) Behavioural light preference after forward- or backward-paired training protocols for wild-type (Canton-S, grey) and orco mutant (orco^1^, magenta) of L2 feeding-stage larvae. ***p<0.001, ****p<0.0001, Mann-Whitney U test. C) Comparison of early-L2 stage larval turning during the test quantified as a behavioural preference index between wild-type (Canton-S, grey) and orco mutants (orco^1^, magenta). Traces show mean ± SEM. D) Behavioural learning index quantified from the turning differences between forward- and backward-trained wild-type (grey) and orco mutant (magenta) L3 larvae. ****p<0.0001, Mann-Whitney U test. E) Left: Sleep profile of adult flies with intact (grey) and cut (blue) antennae at day 3 after antennae removal. Right: Total sleep duration of intact (grey) and cut (blue) flies. **p<0.01, Mann-Whitney U test. F) Same as (E) but sleep profiling performed at day 7 with intact (grey) and cut (teal) flies. “n.s.”p>0.05, Mann-Whitney U test. G) Left: Odour avoidance assay to probe for the presence of olfaction. Right: Comparison of the odour preference of adult flies with intact (grey) and cut (teal) antennae. *p<0.05, two-tailed Student’s t-test. H) Learning index upon mock training (no shock) of adult flies with intact (grey) and cut (teal) antennae. “n.s.”p>0.05, two-tailed Student’s t-test. Boxplots in (A, B, D, E, F, G, H) show median, interquartile ranges and whiskers extend to the most extreme data points. Sample sizes for A) wild-type: n=19, orco mutant: n=20; B) forward n=71, backward n=72, orco mutant: forward n=68, backward n=63 (larvae); C, D) wild-type: n= 120, orco mutant: n=122 (larvae); E) intact: n=42, cut: n=42 (flies); F) intact: n=38, cut: n=37 (flies); G) intact: n=16, cut: n=16 (groups of flies); H) intact: n=18, cut: n=18 (groups of flies).

**Figure S2:**
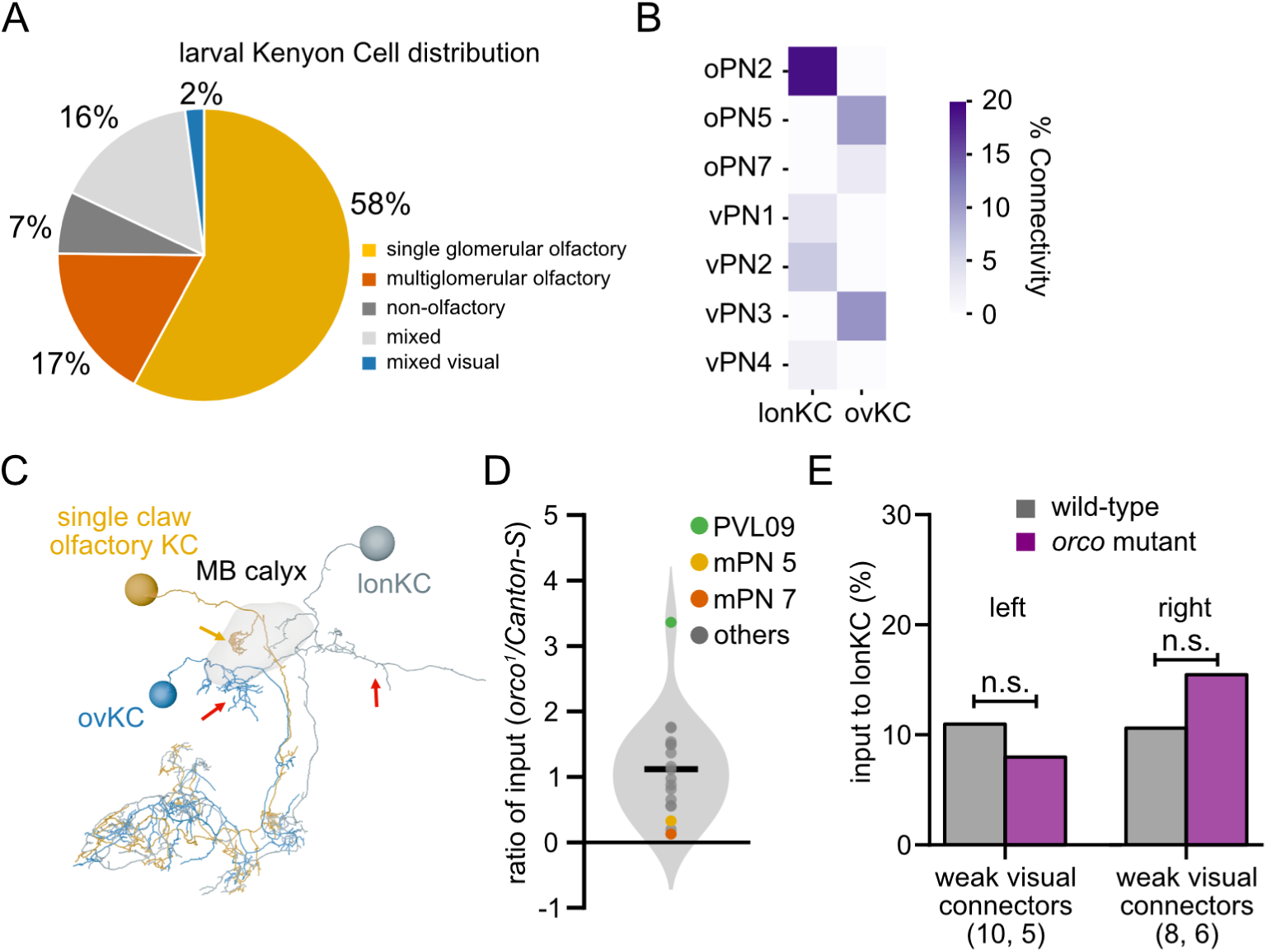
Two distinct Kenyon cells (KCs) receive strong visual input in the larval brain. A) Distribution of all larval KCs based on their input modality. B) Percentage of synaptic input from different presynaptic neurons to lonKC and ovKC in the L1 larval connectome. Both KCs integrate information from visual and multiglomerular olfactory PNs, though only one hemispheric pair of ovKC received visual information in the L1-stage connectome C) The skeletons of ovKC, lonKC and an olfactory KC from the larval connectome. The shaded volume shows the MB calyx. Olfactory KCs show compact dendritic claw structures within the calyx (yellow arrow), whilst both ovKC and lonKC send diffuse dendritic projections outside the calyx (red arrows). D) The ratio between the strength (% connectivity) of homologous connections onto lonKC and ovKC between orco1 and the wild-type volume. Points show the values for each of the 18 homologous input neurons we reconstructed. The shaded area shows the smoothed kernel density estimate for the distribution, with kernel size determined using Scott’s method. Neurons that show significant alterations in both hemispheres are highlighted in colours. E) Comparisons of the total synaptic input that lonKC receives from weak (<3% input) upstream visual neurons in each hemisphere. Upstream neurons were defined as a ‘visual connector’ based on whether they have dendritic arbours and postsynaptic sites within the LON. In addition, weak connections directly from photoreceptors were included in this grouping. The total number of weak connectors in the wild-type and orco mutant volume is shown in brackets (wild-type, orco). The total input from weak visual connectors (<3% input fraction) onto lonKC did not differ significantly between the orco mutant and wild-type volumes. Thereby, while visual excitatory connections onto one multisensory KC were strengthened, the other remained unchanged following loss of olfactory signalling. ns: p>0.05, Chi-square test. The data source for A-C was the previously published L1 larval connectome 35,49.

**Figure S3:**
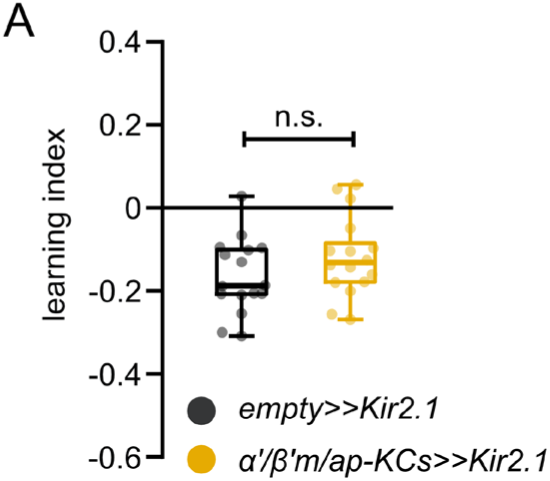
Silencing olfactory KCs does not alter visual learning. A) Visual learning performance of adult flies upon silencing of olfactory KCs (dark yellow). “n.s.”p>0.05, two-tailed Student’s t-test. Boxplots show median, interquartile ranges and whiskers extend to the most extreme data points. Sample size for A) empty>>Kir2.1: n=16, α′β′(ap and m)-KCs>>Kir2.1: n=16.

**Figure S4:**
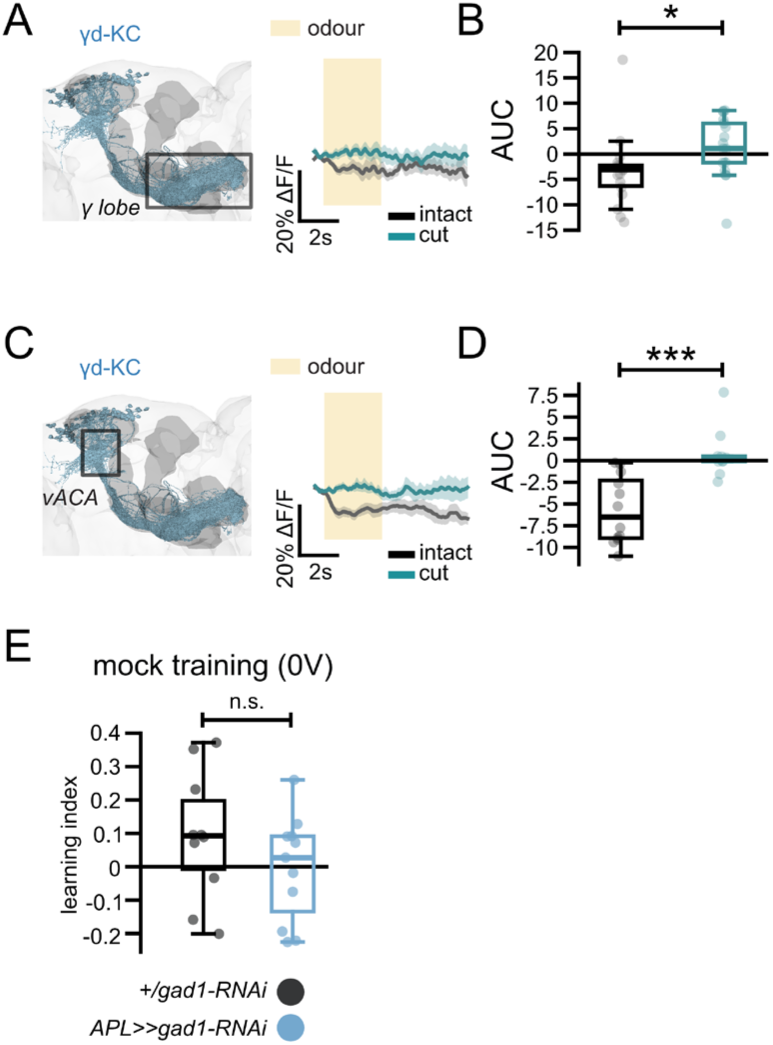
γd-KC responses are suppressed by odours. A) Calcium responses of γd-KCs in the MB lobes upon odour stimulation in flies with intact (black) and cut (teal) antennae. B) Comparison of AUC of the γd-KC calcium responses between intact (black) and cut (teal) flies. C, D) Same as (A, B) but calcium responses of γd-KCs are imaged at their dendrites at the vACA. E) Learning index upon mock training (no shock) of adult flies expressing gad1-RNAi under the control of APL driver (blue) and RNAi controls (grey). Traces show mean ± SEM. “n.s.”p>0.05, *p<0.05, ***p<0.001, two-tailed Student’s t-test. Boxplots in (B, D, E) show median, interquartile ranges and whiskers extend to the most extreme data points, excluding the outliers. Sample sizes for (A, B) intact: n=18, cut: n=17 (flies); (C, D) intact: n=12, cut: n=10 (flies); (E) +/gad1-RNAi: n=10, APL>>gad1-RNAi: n=11 (groups of flies).

## Notes

### Competing Interest Statement

The authors have declared no competing interest.

### Summary of Updates

The Acknowledgements section has been updated.

